# Biotin-operated ligand trapping enables dynamic spatial redistribution of protein and organelle in living cells

**DOI:** 10.64898/2026.10.02.756291

**Authors:** Wyatt C. Beyers, Sarah E. Plutkis, Kathryn R. Butcher, Julianna L. Sun, MaryEllen R. Haas, Mollie S. Hutton, Hark Kyun Kim, Alejandro Chavez, Shondra M. Pruett-Miller, Jason D. Vevea, Luke D. Lavis, Chi-Lun Chang

## Abstract

Controlling protein localization through chemical biology tools is essential for mechanistic interrogations, functional perturbations, and therapeutic development. While existing tools cover a wide range of applications, no single approach simultaneously enables inducible trapping, controlled release, and visualization of a protein of interest (POI). To address this gap, we developed Biotin-Operated Ligand Trapping (BOLT), which combines a HaloTag-fused POI with a streptavidin trap. Desthiobiotin-conjugated Janelia Fluor HaloTag ligands enable POI visualization, trapping, and biotin-operated release. We demonstrate BOLT’s capacity in synchronized secretory cargo release, microtubule-directed organelle movement and hitchhiking, and ectopically localizing nuclear transcription factor. Importantly, pulse-chase BOLT further reveals bidirectional endoplasmic reticulum–Golgi trafficking of an endogenous Golgi enzyme and distinguishes recycled from newly synthesized enzyme populations using distinct HTLs. Altogether, BOLT provides an all-in-one platform to interrogate how subcellular localization regulates protein function across HaloTag-compatible systems.

## Main

Eukaryotic cells partition their cytoplasm into dynamic spatial organizations, such as organelles and cytoskeletal machinery, to support diverse biochemical processes carried out by ∼20,000 protein effectors^1^. Controlling protein localization via synthetic and chemical biology approaches has been proven a powerful avenue to interrogate protein functions and influence cellular fates^2,3^. The classical FK506-binding protein (FKBP) and FKBP-rapamycin binding (FRB) pair, which heterodimerize upon binding to rapamycin and its analogs^4–7^, has been applied to sequester structural components, trafficking machinery, and enzymes to designated subcellular loci for functional interrogation, augmentation, and inhibition both *in vitro* and *in vivo*^8–11^. Paired streptavidin and streptavidin-binding peptide (SBP) have been applied for the Retention Using Selective Hooks (RUSH) system whereby the trafficking of SBP-appended secretory cargos is constitutively halted via trapping with an endoplasmic reticulum (ER)-resident streptavidin-KDEL hook^12^. The addition of biotin frees the SBP-cargos from the hook, leading to a synchronized, anterograde cargo movement, thus enabling quantitative tracking as the cargos traverse the cell’s secretory compartments. While these two systems are widely applicable, the FRB–rapamycin–FKBP ternary complex is practically irreversible within reasonable time frames^5,13^ and streptavidin–SBP binding is virtually constitutive^14,15^, prohibiting inducible interaction.

Recent expansion of chemically induced dimerization systems has partially overcome this limitation. For example, the salicylic acid (SA)-Mediated Binary Association (SAMBA) system, engineered from a tobacco SA receptor, enables rapid SA-mediated heterodimerization to drive receptor signaling and gene expression^16^. Chemically Assisted Tethering of CHimera by Fluorogenic-Induced REcognition (CATCHFIRE) leverages fluorogen-induced complementation of the splitFAST system, a fluorescence complementation system that enables visualization of protein–protein interactions^17^, to alter protein localization and control organelle positioning^18^.

Furthermore, the dimerizing fluorogen in the CATCHFIRE system provides a high-precision readout of the interaction foci. Even though inducible trapping in SAMBA and CATCHFIRE are reversible, these systems lack a visualization component to monitor the protein of interest (POI) during the dynamic trap-release cycles.

The HaloTag is a genetically encoded self-labeling enzyme tag that can covalently react with a HaloTag ligand (HTL) conjugated with a large selection of small molecules, including Janelia Fluor (JF) dyes^19,20^. Given its brightness, photostability, fluorogenicity, and imaging capacity across a wide range of the light spectrum, the HaloTag has been broadly adopted as a fluorescent reporter for a variety of imaging modalities, including super-resolution imaging^20^. In addition to fluorescent HTLs for imaging, a variety of nonfluorescent ligands exist with unique functional groups such as a biotin, for biochemical enrichment of HaloTag-appended proteins ^21^. Recently described multifunctional HTLs have further combined a JF dye and a biotin moiety in the same molecule (biotin-JF-HTL), enabling the visualization and enrichment of HaloTag-appended proteins and organelles via the same molecular handle^22^. Conceptually, these biotin-JF-HTLs are capable of simultaneous labeling and inducible dimerization upon addition of a ligand with co-expression a streptavidin-tagged trap, analogous to FRB-FKBP system with a built-in fluorescent tag for POI. Although the interaction between biotin and streptavidin with a dissociation constant (K*_d_*) in the femtomolar range (6.5×10^-14^M) is essentially irreversible under physiological conditions, the binding of streptavidin and desthiobiotin (DTB), with a K*_d_* _o_f 6.6×10^-10^M, is readily reversible upon the addition of biotin^23–25^. Therefore, a DTB-conjugated JF-HTL that covalently links to a HaloTag and reversibly binds to streptavidin could overcome the technical limitations of all existing dimerization systems; this would enable precise detection and rapid, reversible localization control of a POI for causal mechanistic studies. This new, multifunctional HTL would be broadly impactful, particularly because HaloTag is already widely used across diverse biological systems.

Here, we describe the development of biotin-operated ligand trapping (BOLT), a system using a new class of multi-functional HTL consisting of a JF dye for visualization and DTB for reversible trapping to streptavidin. We validate that these new DTB-JF-HTLs are intracellularly available and subjected to biotin operated, reversible interaction with streptavidin *in vitro* and within cells. Functionally, we demonstrate that the BOLT system, when applied to synchronized secretory cargo release, is comparable to the benchmark RUSH approach. In addition to controlling protein trafficking, BOLT is capable of directing mitochondria distribution via microtubule-mediated transport, revealing peroxisome, but not lipid droplet (LD), hitchhiking on moving mitochondria, suggesting different interactions between mitochondria and other organelles.

Furthermore, BOLT can regulate transcription factors nuclear import, providing proof-of-concept for its potential application in regulating transcriptional programs. Importantly, BOLT is readily applicable to the control of endogenous HaloTag-fused proteins, revealing bidirectional trafficking between the ER and Golgi of an endogenous Golgi enzyme, MAN2A1, as well as regulating nuclear import of endogenous p53 transcription factor. Altogether, our data demonstrated that BOLT is an all-in-one, generalizable system that provides dynamic trap-release and simultaneous visualization of POI via HaloTag and a multifunctional HTL.

## Results

### Design and synthesis of DTB-JF-HTL for implementing BOLT

In order to develop the BOLT system for dynamic, reversible trapping and release and simultaneous visualization of POI based on the HaloTag and streptavidin pairing, we first generated a multifunctional HTL containing DTB, to allow the reversible release of HaloTag-fused proteins or organelles upon biotin addition, and a JF dye, for imaging purposes. Previous work demonstrated biotin-JF molecules based on Si rhodamine dyes could efficiently permeate cells^22^. We chose two dyes, JF_646 a_nd JF_635,_ due to their bright, far-red fluorescence^26^, to include in our multifunction HTL design. JF_635-_HTL has the added benefit of being highly fluorogenic^27^, which meant the DTB-JF_635-_HTL molecule should exhibit a substantial increase in fluorescence upon reaction with the HaloTag. Following a similar strategy used for the biotin-JF-HTL compounds^22^, we synthesized two new molecules: DTB-JF_646-_HTL and DTB-JF_635-_HTL, where the DTB moiety is attached to one of the azetidine rings on the JF dyes via a short polyethylene glycol (PEG) linker (Fig. 1A).

**Figure 1.**
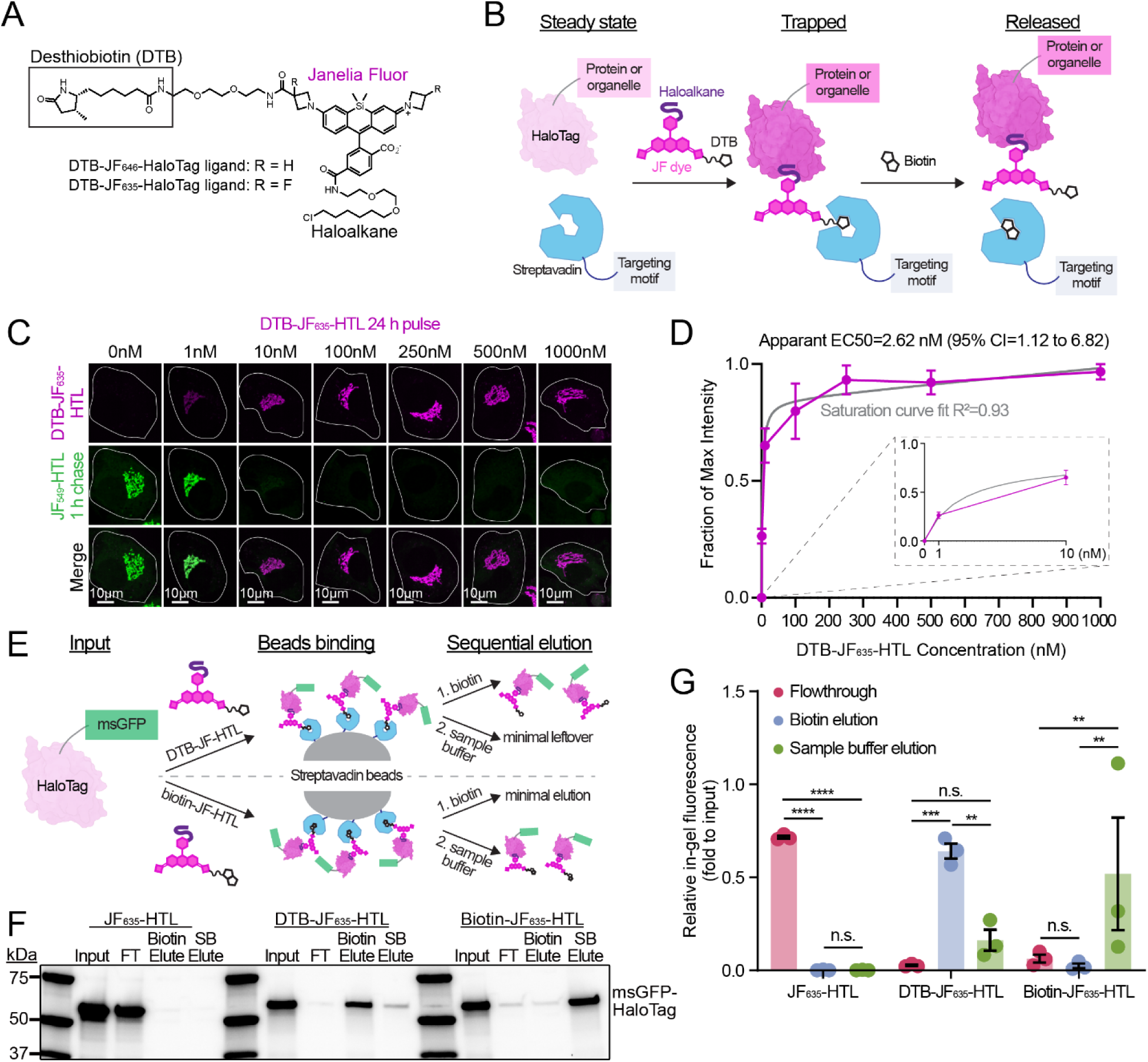
Design, generation, and validation of desthiobiotin-Janelia Fluor-HaloTag ligands (DTB-JF-HTLs) for the Biotin-Operated Ligand Trapping (BOLT) system. (A) Chemical structure of DTB-JF-HTL. The HTLs contain a DTB moiety for capture by streptavidin, a far-red JF dye for fluorescence, and a haloalkane linker to react with the HaloTag. (B) A diagram illustrating the BOLT system. BOLT utilizes a HaloTag-fused protein of interest and a streptavidin with a targeting motif. DTB-JF-HTL then covalently labels HaloTag and induces an interaction with streptavidin to trap it. The addition of biotin will reverse the interaction, thus releasing the HaloTag. The HaloTag illustration was made using Biorender based on HaloTag’s crystal structure (PDB: 5Y2X). (C) Confocal maximum intensity projections of HaloTag knock-in RPE1 cells that were incubated with varying concentrations of DTB-JF_635-_HTL for 24 h, fixed, and stained with 1 µM JF_549-_HTL to label leftover HaloTag. (D) Saturation curve of DTB-JF_635-_HTL mean fluorescence intensity (background subtracted) corresponding to images in (C). Experimental mean intensity values are min-max normalized. The plot points and error bars indicate mean ± S.E.M. The saturation curve fitting was performed assuming one site binding setting in Graphpad Prism. (E) A diagram depicting the streptavidin pulldown assay to assess of biotin-JF-HTL’s and DTB-JF-HTL’s ability to bind to streptavidin-coated magnetic beads, and if the molecules can be released with biotin. (F) In-gel fluorescence SDS-PAGE gel showing the results of a streptavidin pulldown assay comparing the input, flowthrough (FT), first biotin elution, and second elution using sample buffer (SB) with heat for JF_635-_HTL, DTB-JF_635-_HTL, and biotin-JF-HTL. (G) Quantification of in-gel fluorescence SDS-PAGE gels as shown in (F). The in-gel fluorescence was normalized to the input; quantification occurred across three independent experiments. Data is presented as mean ± S.E.M. Significance was tested using a two-way ANOVA with pairwise multiple comparisons; n.s., not significant; \*\**P* < 0.01; \*\*\**P* < 0.001; \*\*\*\**P* < 0.0001.

These newly generated multifunctional HTLs fulfill the missing component for the BOLT system, which concurrently (1) labels and conjugates the HaloTag-fused POI and (2) interacts with the streptavidin trap targeted to designated subcellular foci (Fig. 1B). The addition of biotin displaces the interaction between streptavidin and the DTB moiety, enabling a rapid release of the DTB-JF-HTL–appended POI to restore the POI’s localization and function. We validate and demonstrate the feasibility of BOLT in the secretory pathway, organelle position, and nuclear import of transcription factors.

### Validation of intracellular availability and reversibility of DTB-JF-HTLs

We first examined the intracellular availability of DTB-JF_635-_HTL and DTB-JF_646-_HTL, as this is critical to implement BOLT, by pulsing HEK293 cells overexpressing a cytosolic HaloTag with each HTL for 2 h, followed by chasing with a highly permeable ligand, JF_549-_HTL, to label any remaining unlabeled HaloTag. The in-gel fluorescence assay showed that, despite less overall HaloTag labeling than would occur with a standard JF-HTL lacking functional moiety, the DTB-JF-HTLs successfully labeled intracellular HaloTags to levels comparable to those observed with biotin-JF-HTLs (Extended Data 1A–D). We then sought to determine if a longer incubation period with a DTB-JF-HTL could fully label an endogenously HaloTag-fused Golgi enzyme, MAN2A1, in RPE1 cells, as we intended to use BOLT to dissect Golgi enzyme redistribution (see Fig. 3). These cells were incubated with a range of DTB-JF_635-_HTL concentrations for 24 h, fixed, and stained with JF_549-_HTL for 1 h to assess DTB-JF_635-_HTL labeling efficiency (Fig. 1C, 1D). At a concentration of 100 nM or greater, DTB-JF_635-_HTL labeled almost all of the endogenous MAN2A1-HaloTag with a half maximal effective concentration (EC_50)_ of 2.62 nM (95% CI: 1.12–6.82 nM), while 1 nM of DTB-JF_635-_HTL only labeled a small population of MAN2A1-HaloTag. This suggests a range of 100–500 nM is suitable to achieve optimal DTB-JF-HTL labeling in situations when a 24-h incubation is possible.

We next assessed the reversibility of DTB-JF-HTL trapping using a streptavidin-bead pulldown assay (Fig. 1E). Recombinant HaloTag proteins incubated with DTB-JF_635-_HTL or biotin-JF_635-_HTL were efficiently enriched onto the streptavidin beads while HaloTag proteins incubated with JF_635-_HTL as a negative control were only present in the flowthrough (Fig. 1F, 1G). To assess reversibility of DTB-JF_635-_HTL or biotin-JF_635-_HTL, we performed a sequential elution with biotin followed by sample buffer with heat, and found that the addition of biotin was sufficient to elute the majority of DTB-JF_635-_HTL labeled HaloTag while biotin-JF_635-_HTL-conjugated proteins were only eluted when we used sample buffer and applied heat. Similar results were observed when we compared JF_646-_HTL, DTB-JF_646-_HTL, and biotin-JF_646-_HTL (Extended Data 2A, 2B). We then tested the binding capabilities for these ligands in the cytosol of cells to determine if they would perform similarly in a more physiologically relevant context. Within RPE1 cells, a cytosolic streptavidin-BFP-nuclear export signal (NES) trapped proteins labeled with biotin-JF_646-_HTL and DTB-JF_646-_HTL, but not JF_646-_HTL. Biotin addition reversed the streptavidin–DTB-JF_646-_HTL interaction, while biotin-JF_646-_HTL signal persisted within cells even after adding biotin, indicating DTB-JF-HTL binding is reversible (Extended Data 2C, 2D). These findings established DTB-JF-HTL as optimal ligands for use with BOLT, as they induced a robust but reversible HaloTag–streptavidin interaction both *in vitro* and in cells.

### Applying BOLT to monitor ER-to-Golgi protein trafficking

To further test BOLT’s feasibility within a biological context, we adapted BOLT for synchronized secretory cargo release with an ER-resident streptavidin-KDEL trap and a synthetic secretory cargo containing mouse MAN2A1’s N-terminal (aa 1–102), mNeonGreen (mNG) as an independent localization readout, and a HaloTag for BOLT implementation, hereafter referred to as Man2 N’-mNG-HaloTag (illustrated in Fig. 2A). In RPE1 cells, JF_635-_HTL-labeled Man2 N’-mNG-HaloTag trafficked freely to the Golgi as they colocalized with a Golgi marker SiT-FusionRed at steady state, while overnight DTB-JF_635-_HTL incubation resulted in cargo accumulation in the ER with unaffected SiT-FusionRed distribution (Fig. 2B), reflecting inducible trapping of cargo via BOLT by DTB-JF_635-_HTL while ER-to-Golgi trafficking remain unaffected.

**Figure 2.**
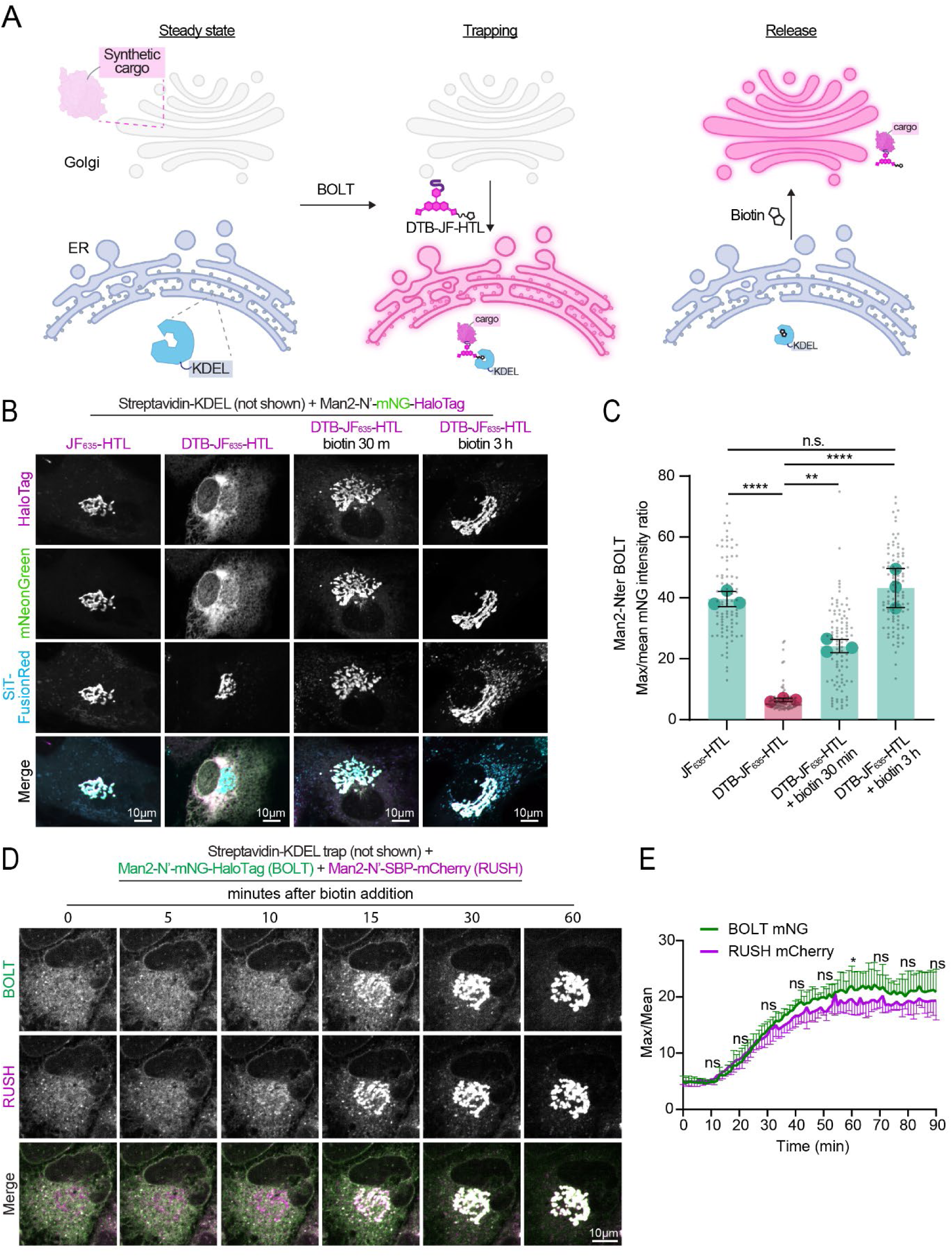
Applying BOLT to monitor synchronized endoplasmic reticulum (ER)-to-Golgi protein trafficking. (**A**) A diagram depicting how the BOLT system traps and releases synthetic secretory cargos using an ER-localized trap, streptavidin-KDEL. Secretory cargo with a HaloTag is trapped in the ER by adding desthiobiotin-Janelia Fluor-HaloTag ligand (DTB-JF-HTL) and released by adding biotin. (**B**) Confocal maximum intensity projections of RPE1 cells transduced with streptavidin-KDEL IRES Man2-N’-HA-mNeonGreen (mNG)-HaloTag and SiT-FusionRed lentiviral particles and incubated with 100 nM JF_635-_HTL or DTB-JF_635-_HTL overnight, then released upon incubation with biotin (80 µM) for 30 or 180 min. (**C**) Max-to-mean intensity ratio per cell for mNG fluorescence of images shown in (B). Bars depict the mean ± S.D. from three independent experiments (shown as large points), while small points represent individual (n=86–101) cells per condition. Statistics were performed using a one-way ANOVA with *post hoc* multiple comparisons; n.s., not significant; *\*\*P <* 0.01; \*\*\*\**P* < 0.0001. (**D**) Confocal 2D slices of RPE1 cells transduced with lentiviral particles encoding streptavidin-KDEL IRES Man2-N’-HA-mNG-HaloTag and streptavidin-KDEL IRES Man2-N’-SBP-mCherry, incubated with DTB-JF_635-_HTL overnight to trap, and released with 80 µM biotin immediately after the first frame to assess trafficking dynamics of BOLT cargo compared to RUSH cargo. Man2-N’ BOLT images are showing the mNeonGreen signal. (**E**) Quantification of release dynamics for timelapses shown in (D). Traces indicate the max-to-mean intensity ratio per cell over time to quantify ER-to-Golgi trafficking, shown as mean ± S.D. from the independent experiments. Paired *t*-tests were performed for various timepoints to evaluate differences in dynamics between the BOLT and RUSH cargoes (n=6–14 cells per experiment); n.s., not significant; *\*P <* 0.05.

Addition of biotin rapidly reversed the cargo retention leading to a synchronous ER-to-Golgi Man2 N’-mNG-HaloTag trafficking within 30 min and retained in the Golgi after a 3-h biotin incubation (Fig. 2B). Throughout these experiments, we observed protein redistribution from ER to Golgi can be reliably quantified by measuring the maximum pixel intensity within region of interest (ROI) within a cell because the diffuse fluorescence intensity signal through the entire ER becomes tightly concentrated in one region (the central Golgi), greatly increasing maximum intensity in the cell. Therefore, we measured maximum intensity normalized to mean (hereafter referred to as max-to-mean) intensity ratio, which measures the tight clustering of signal which causes heightened maximum pixel intensity per cell while normalizing to mean intensity per cell, to assess ER-to-Golgi trafficking (Fig. 2C). Similar trap-release cycle was also observed with another synthetic cargo engineered from TNFα, a secreted protein with a transmembrane domain (Extended Data 3A, 3B).

As RUSH is a well-established system for synchronized cargo release and shares similar trapping and release mechanisms with BOLT, we sought to determine if BOLT performs comparably to the benchmark RUSH system. In RPE1 cells expressing streptavidin-KDEL, we observed ER retention of both Man2-N’-SBP-mCherry (cargo for RUSH) and Man2-N’-mNG-HaloTag (cargo for BOLT) in the presence of DTB-JF_635-_HTL (Fig. 2D). Following the addition of biotin, ER-to-Golgi trafficking of both cargos occurred within a comparable timeframe, with prominent signal visible in the Golgi 15 min after biotin-induced release (Fig. 2D). Max-to-mean intensity analysis revealed no significant difference in trafficking rate; however, at the 60-min mark, the BOLT cargo had modest increase in max-to-mean intensity ratio, suggesting a slight improvement in trafficking capacity over to the RUSH system (Fig. 2E).

A major impetus for developing BOLT was its potential to manipulate endogenous proteins with a HaloTag fusion via CRISPR knock-in. In theory, cell lines originally generated for steady state imaging could be repurposed for functional trafficking assays without the need to knock in an additional tag. We assessed whether we could use BOLT to monitor ER-to-Golgi trafficking of the endogenous Golgi resident enzyme, MAN2A1, homozygously tagged with a HaloTag via CRISPR knock-in in RPE1 cells. We first confirmed endogenous MAN2A1-HaloTag labeled with JFX_646-_HTL colocalized with the Golgi immunostained using Golgin97 antibody in knock-in RPE1 cells expressing streptavidin-KDEL. Following overnight incubation with DTB-JF_646-_HTL, endogenous MAN2A1-HaloTag was retained in the ER and the Golgi was intact as shown by the Golgin97 immunostaining. The trapped MAN2A1 was subsequently released via addition of biotin and was trafficked to the Golgi (Extended Data 3C, 3D). Together, these results demonstrate that BOLT is capable of visualizing ER-to-Golgi protein trafficking at comparable kinetics to the benchmark RUSH system and is applicable for use with endogenous secretory cargos fused with a HaloTag. In addition, BOLT allows constitutive cargo trafficking at steady state and inducible trapping upon addition of a DTB-JF-HTL, which should minimize a cell’s stress response to prolonged accumulation of overexpressed cargo in the ER, as occurs in the RUSH system.

### BOLT reveals bidirectional ER-Golgi trafficking of endogenous Golgi resident enzyme

Multiple reports have suggested that Golgi resident proteins can constitutively recycle from Golgi back to the ER^28–30^. This phenomenon can be visualized by capturing synthetic, overexpressed Golgi-resident reporters in the ER using a trapping system based on FRB-FKBP over a course of several hours following the addition of rapamycin and its analog^28^. However, whether endogenous Golgi enzymes undergo this ER recycling and whether recycled enzymes remain capable of ER-to-Golgi trafficking remain unclear. To test if endogenous MAN2A1-HaloTag can redistribute to the ER, we devised a pulse-chase BOLT assay, where the knock-in cells were pulse labeled with DTB-JF_646-_HTL in the absence of streptavidin-KDEL expression (Fig. 3A). Without a trap, endogenous MAN2A1-HaloTag was localized to the Golgi with negligible ER distribution. Following doxycycline-induced streptavidin-KDL expression in the absence of DTB-JF_646-_HTL for 6 h, the existing MAN2A1-HaloTag-DTB-JF_646 w_as accumulated in the ER, suggesting a robust, albeit slow, Golgi-to-ER recycling of MAN2A1-HaloTag over time (Fig. 3B, 3C). In addition, we quenched any unlabeled MAN2A1-HaloTag with 7-bromo-1-heptanol and then chase labeled the cells with JFX_549 d_uring doxycycline incubation to label the newly synthesized population of MAN2A1-HaloTag and found mainly Golgi distribution of MAN2A1-HaloTag-JFX_549 (_Fig. 3B, 3C), indicating functional ER-to-Golgi trafficking. Intriguingly, biotin addition also led to the rapid redistribution of MAN2A1-HaloTag-DTB-JF_646 t_o the Golgi within 45 min, completing a full round of Golgi-to-ER recycling and subsequent ER-to-Golgi trafficking (Fig. 3D, 3E). In negative control experiments using JFX_646-_HTL instead of DTB-JF_646-_HTL, the POI failed to redistribute from Golgi to ER (Extended Data 4A, 4B). Altogether, these results indicate the pulse-chase BOLT assay will facilitate the assessment of bidirectional ER-Golgi trafficking of endogenous cargo and the characterization of the recycling population of enzymes from those that are newly synthesized by using distinct HTLs.

**Figure 3.**
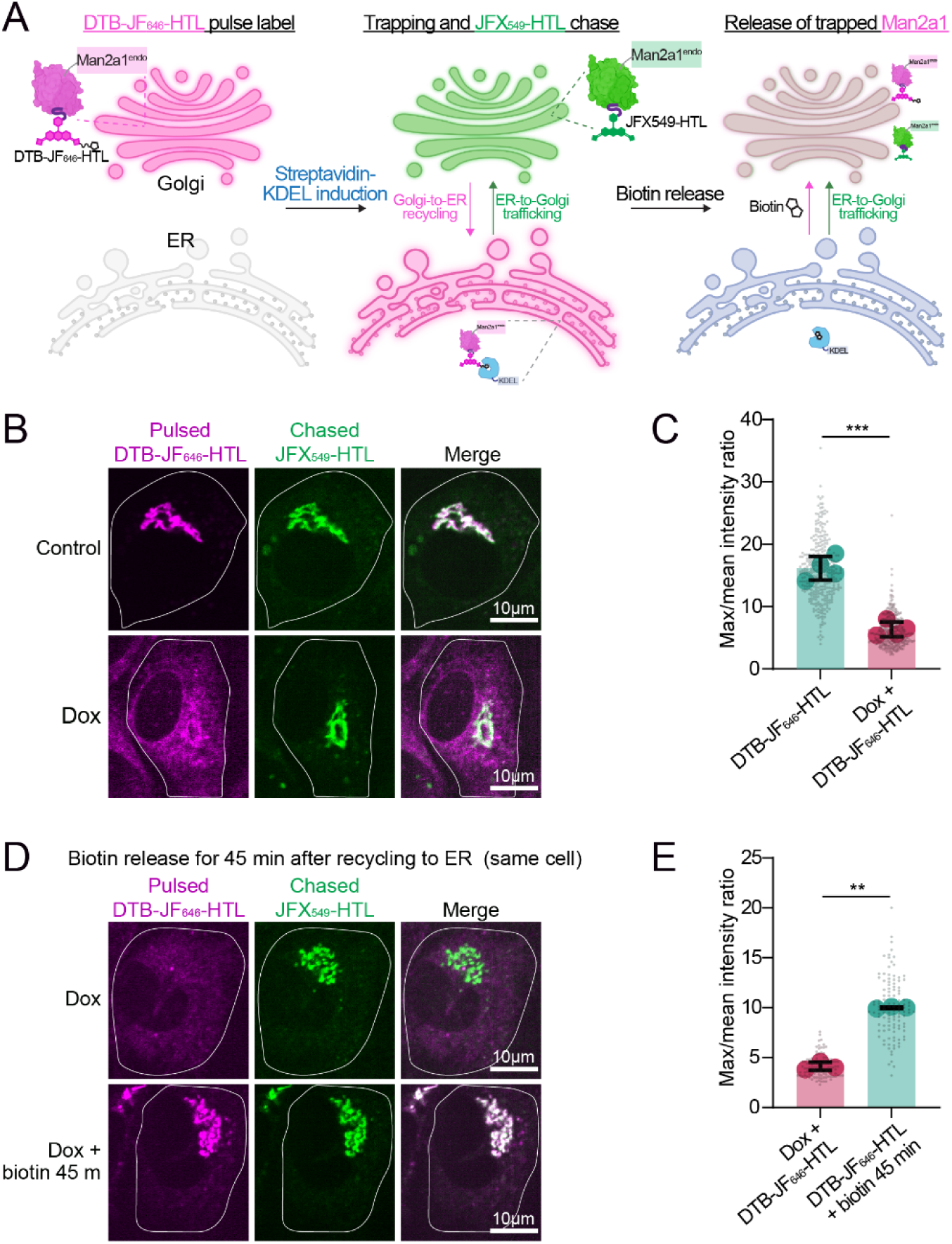
BOLT reveals bidirectional endoplasmic reticulum (ER)-to-Golgi trafficking of endogenous Golgi resident enzyme. (**A**) A diagram illustrating the pulse-chase BOLT assay for endogenous MAN2A1-HaloTag in RPE1 cells. MAN2A1-HaloTag was pulse labeled overnight with desthiobiotin-Janelia Fluor_646-_HaloTag ligand (DTB-JF_646-_HTL) to label a steady-state Golgi population. After washing out the dye and quenching with 7-bromo-heptanol, cells were incubated with doxycycline (dox) and JFX_549-_HT to induce streptavidin-KDEL expression and chase-label newly synthesized, Golgi-bound MAN2A1-HaloTag as it was formed. After streptavidin-KDEL was induced, the pre-existing recycling population of MAN2A1-HaloTag-DTB-JF_646-_HTL was trapped in the ER. The addition of biotin released the trapped MAN2A1-HaloTag-DTB-JF_646-_HTL population, which could traffic back to the Golgi, completing a full round of recycling. (**B**) Confocal maximum intensity projection (MIPs) of MAN2A1-HaloTag knock-in RPE1 cells that were pulsed overnight with 100 nM DTB-JF_646-_HTL and chased with 100 nM JFX549-HTL ± 1 µg/mL doxycycline for 6 h to induce streptavidin-KDEL expression. (**C**) Max-to-mean intensity ratios of images from (B), graph depicts mean ± S.D. from four independent experiments (shown as large, colored points). Small, grey points indicate individual (n=334–360) cells per condition. A Welch’s *t*-test was performed to test statistical significance; \*\*\**P* < 0.001. (**D**) Confocal MIPs of MAN2A1-HaloTag knock-in RPE1 cells that were pulsed overnight with 100 nM DTB-JF_646-_HTL and chased with 100 nM JFX549-HTL with 1 µg/mL doxycycline for 6 h to induce streptavidin-KDEL expression, before 45-min incubation with biotin (80 µM) to monitor MAN2A1-HaloTag-DTB-JF_646-_HTL’s ability to traffic from the ER back to the Golgi, completing a full round of recycling. (**E**) Max-to-mean intensity ratios of images from (D); graph depicts mean ± S.D. from three independent experiments (shown as large, colored points). Small, grey points indicate (n=108–116) individual cells per treatment. A Welch’s *t*-test was performed to test statistical significance; \*\**P* < 0.01.

### BOLT mediates microtubule-directed organelle movement and hitchhiking

After we used BOLT to successfully induce the trapping and release of our POI within a cell’s secretory compartments, we next sought to apply BOLT to manipulate the distribution of entire organelles. Several organelles (e.g., mitochondria, peroxisomes) utilize motor proteins to traffic progressively along microtubule tracks via kinesin-mediated plus-end trafficking toward the cell’s periphery or via dynein-directed minus-end movement towards the cell’s center^31^. Dynein is recruited and activated by organelle-specific adapter proteins, such as BICD2, to initiate organelle trafficking. Several groups have demonstrated chemically induced BICD2 recruitment to peroxisomes promoted peroxisome accumulation in the center of cells^32–34^. Therefore, we exogenously expressed streptavidin-fused BICD2 to test if BOLT could be employed to control movement of HaloTag-labeled organelles, starting with TOM20-HaloTag–labeled mitochondria (Fig. 4A). Within minutes of adding DTB-JF_635-_HTL, mitochondria started moving toward the cell center, forming a tight central cluster by 30 min (Fig. 4B). We quantified this movement by measuring the max-to-mean intensity ratio (Fig. 4C). After 30 min of clustering, we added biotin, which caused BICD2-streptavidin release and restored native anterograde trafficking, and the mitochondria to redispersed. During BOLT-induced mitochondria minus-end trafficking, we noted the mitochondria became fragmented during transit (Fig. 4B insets, 4D, 4E). These experiments demonstrate BOLT can acutely influence mitochondria trafficking along microtubules coinciding with changes in mitochondrial morphology.

**Figure 4.**
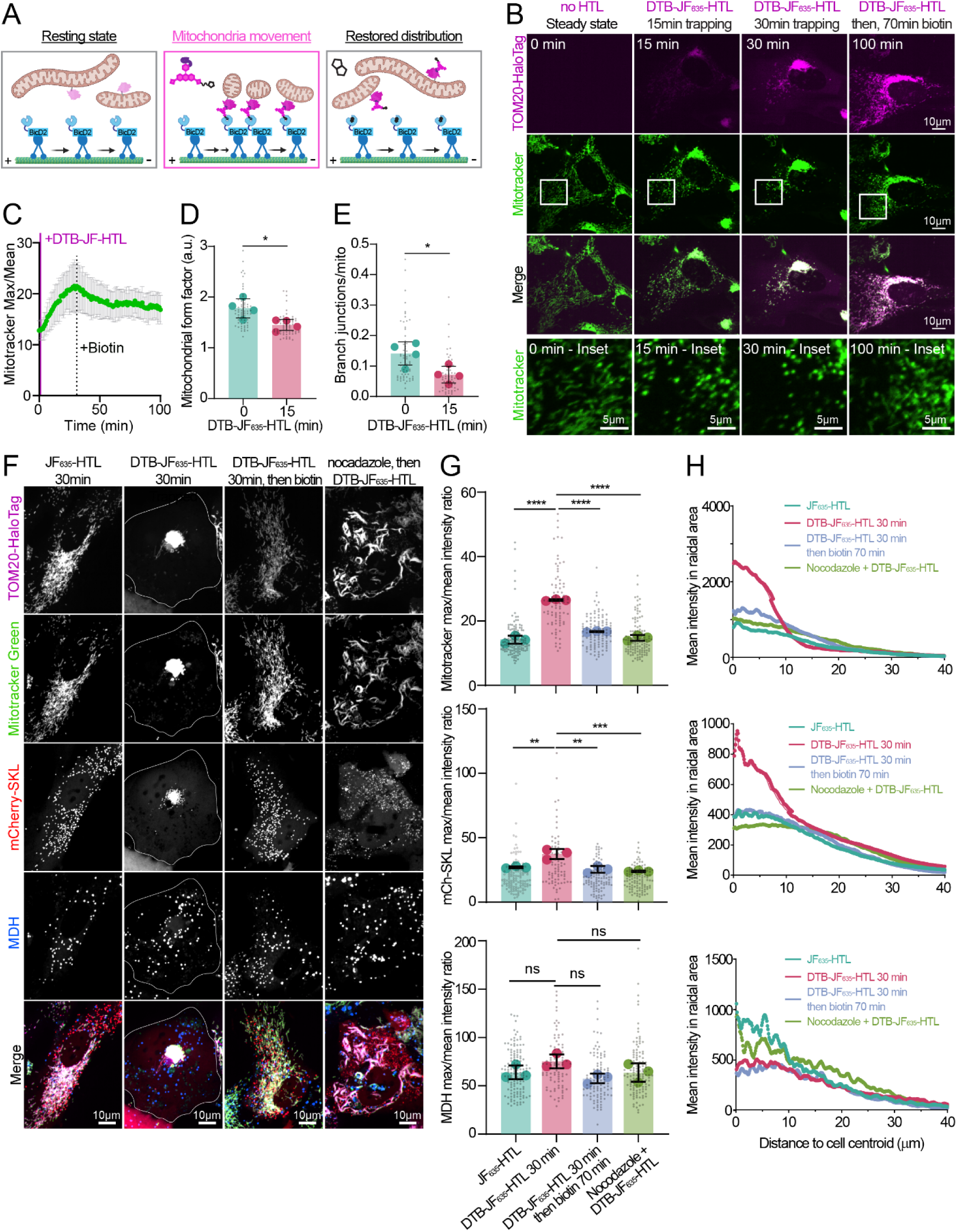
BOLT mediates microtubule-directed organelle movement and hitchhiking. (**A**) Diagram depicting BICD2-streptavidin BOLT assay. At steady state, BICD2-streptavidin is not coupled to mitochondria. Desthiobiotin-Janelia Fluor_635-_HaloTag ligand (DTB-JF_635-_HTL) simultaneously labels Tom20-HaloTag and induces an interaction with BICD2-streptavidin, initiating minus-end directed movement on microtubules via the dynein motor complex. The addition of biotin releases TOM20-HaloTag-DTB-JF_635-_HTL from BICD2-streptavidin, allowing normal mitochondria distribution to be restored. (**B**) Confocal 2D images from timelapse imaging for RPE1 cells that were transduced with flag-BicD2-streptavidin and TOM20-HaloTag, and stained with 100 nM MitoTracker Green FM. One-hundred nanomolar DTB-JF_635-_HTL was added immediately after the first frame of the timelapse and 80 µM biotin was added after 30 min of trapping to test reversibility. Insets show areas with tubular mitochondria at the start of the timelapse that appear to become fragmented during induced trafficking. (**C**) Quantification of the mitochondria trafficking phenotype shown in (B). MitoTracker Green max-to-mean intensity ratio per cell over the course of timelapse imaging, with points and error bars depicting the mean ± S.D. from three independent experiments with n=30 cells analyzed. The vertical magenta line indicates the frame where DTB-JF_635-_HTL was added, while the vertical black dotted line indicates the frame where biotin was added to induce release. (**D**) Mitochondrial form factor determined Mitochondria Analyzer for images in (B) measured at the first frame (time=0) and 15 min after adding DTB-JF_635-_HTL, which is when mitochondria were in transit to the cell center. The graph depicts the mean ± S.D. from four independent experiments (depicted by large, colored points) and n=61–63 cells/timepoint (depicted as small, grey points). A paired *t*-test was performed to test for significance; \**P* < 0.05. (**E**) Mitochondrial branch junctions per mitochondrion for images in (B) measured using Mitochondria Analyzer at the first frame (time=0) and 15 min after adding DTB-JF_635-_HTL, when mitochondria were in transit to the cell center. Mean ± S.D. from four independent experiments (depicted by large, colored points) and n=61–63 cells/timepoint (depicted as small, grey points) are displayed. A paired *t*-test was performed to test for significance; \**P* < 0.05. (**F**) Confocal maximum intensity projection of RPE1 cells that were transduced with lentiviral particles encoding TOM20-HaloTag, mCherry-SKL, and flag-BicD2-streptavidin, stained with MitoTracker Green and monodansylpentane (MDH), and then incubated with 100 nM JF_635-_HTL (negative control), 100 nM DTB-JF_635-_HTL (BICD2-streptavidin recruited to mitochondria), 100 nM DTB-JF_635-_HTL with 80 µM biotin (for release), or 100 nM DTB-JF_635-_HTL after nocodazole pre-treatment. (**G**) Max-to-mean intensity ratio per cell of images shown in (E). Graphs indicate the mean ± S.D. from three independent experiments (depicted as large, colored points) and n=74–134 cells/condition (depicted as small, grey points). Analysis for MitoTracker green (top), mCherry-SKL (middle), and MDH (bottom) channels are shown as separate graphs. A one-way ANOVA with *post hoc* multiple comparisons was performed to test for significance; ns, no significance; \*\**P* < 0.01; \*\*\**P* < 0.001; \*\*\*\**P* < 0.0001. (**H**) Radial distribution analysis, corresponding to images in (F), depicting the distribution of mean fluorescence intensity in radial bins starting from the cell centroid and moving outward in 0.1-µm increments. The distributions shown depict the mean profile of fluorescence intensity distribution from three independent experiments with n=10 cells/experiment. Profiles were generated using the Radial Profile Angle plugin in Fiji.

Mitochondria are often tethered with other organelles, such as peroxisomes and lipid droplets (LDs), at contact sites to facilitate fatty acid metabolism^35–37^. We wanted to know if these physical interactions would be retained during BOLT-induced mitochondria trafficking, that is, if peroxisomes and LDs would “hitchhike” alongside the mitochondria as they migrated to the center of the cell. To this end, we visualized peroxisomes using mCherry-SKL and LDs using monodansylpentane and found that peroxisomes, but not LDs, moved with mitochondria toward the center of the cell within a similar time frame (Fig. 4F). Clustering of mitochondria and peroxisomes could be reversed after adding biotin to release BICD2-streptavidin, allowing anterograde trafficking to take over, suggesting a robust mitochondria–peroxisome interaction, which was validated with max-to-mean intensity ratio quantification (Fig. 4G). Fluorescence intensity radial distribution also showed changes in mitochondrial and peroxisomal (but not LD) distribution at the 30-min mark of the BOLT assay (Fig. 4H). Pretreating cells with nocodazole to depolymerize microtubules prevented mitochondria from trafficking towards the cell center (Fig. 4F-H), confirming the BICD2-streptavidin BOLT assay is a microtubule-dependent process.

Having established peroxisomes can traffic alongside mitochondria during BOLT-mediated minus-end trafficking, we next investigated if mitochondria could follow peroxisomes when BICD2-streptavidin is recruited to peroxisomes via PMP34-HaloTag. In RPE1 cells co-expressing PMP34-HaloTag and BICD2-stretavidin, the addition of DTB-JF_635-_HTL led to the rapid accumulation of peroxisomes in the cell center. However, mitochondria labeled with Mitotracker green did not redistribute with peroxisomes (Extended Data 5A, 5B, 5C), suggesting the mitochondria–peroxisome interaction was not maintained during peroxisome trafficking. We speculate that the relative size difference between mitochondria and peroxisome may provide an explanation. While smaller peroxisomes can easily hitchhike on mitochondria moving along microtubules, moving peroxisomes may not be able to maintain a strong enough contact to pull a whole mitochondrion along with it. Together, these experiments establish BICD2-streptavidin BOLT as a new tool for inducing and reversing organelle trafficking to monitor organelle dynamics, morphological changes, and interactions with other organelles that occur during microtubule trafficking.

### BOLT-mediated cytosol trapping prevents protein nuclear import

Protein nuclear import and export is important for many cellular functions, including cell cycle and transcriptional controls. We wondered if BOLT could be engineered to manipulate nuclear proteins by trapping them in the cytosol and releasing them with biotin to monitor their nuclear import. We engineered a lentiviral construct encoding doxycycline inducible HaloTag-ATF4 as a model nuclear protein which contains a nuclear import signal to facilitate nuclear translocation, and we sought to test if BOLT could robustly trap it in the cytosol. RPE1 cells were transduced with a cytosolic trap consisting of streptavidin-tagBFP2-NES and doxycycline inducible HaloTag-ATF4 lentiviral particles. Without dox induction, negligible ATF4 expression was detected (Fig.5A). With dox induction and a JF_635-_HTL, HaloTag-ATF4 was readily detected in the nucleus (Fig. 5A). With dox induction and DTB-JF_635-_HTL, HaloTag-ATF4 was successfully trapped in the cytosol, severely losing its nuclear enrichment (Fig. 5A, 5B). Finally, biotin reversal released HaloTag-ATF4 to translocate into the nucleus (Fig. 5A, 5B).

**Figure 5.**
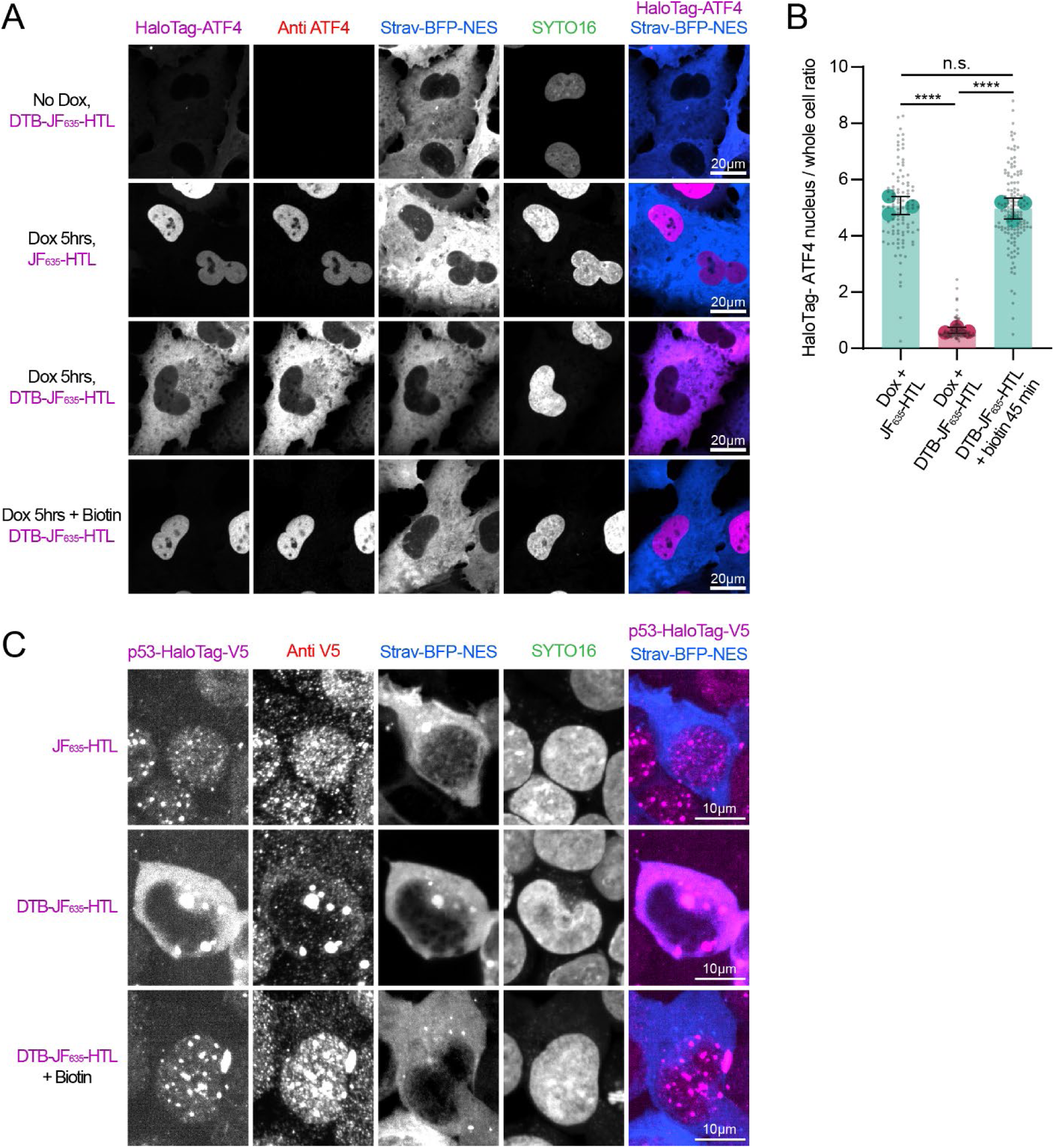
BOLT can control protein nuclear import. (**A**) Confocal maximum intensity projections of RPE1 cells transduced with lentiviruses encoding streptavidin-mtagBFP2-NES and doxycycline-inducible HaloTag-ATF4. They were incubated ± 1 µg/mL doxycycline (to induce HaloTag-ATF4 expression) with desthiobiotin-JF_635-_HaloTag ligand (DTB-JF_635-_HTL; to trap HaloTag-ATF4 in the cytosol) or JF_635-_HTL (negative control) for 5 h. A release condition with doxycycline, DTB-JF_635-_HTL, and 80 µM biotin for the last 3 h of the 5-h induction is also shown to test release. At 5 h after doxycycline incubation, cells were fixed with 4% PFA and immunostained with anti-ATF4 antibody with anti-rabbit Alexa Fluor 555 secondary antibody as an independent marker for ATF4, and stained with SYTO16 DNA dye. (**B**) Quantification showing nuclear enrichment for images shown in (A). The graph depicts the ratio of ATF4 fluorescence intensity in the nucleus to ATF4 fluorescence intensity in the whole cell, with the bars indicating the mean ± S.D. for three independent experiments (shown as large, colored points) and n=103–137 cells per condition (depicted as small, grey points). A one-way ANOVA with *post hoc* Tukey multiple comparisons was used to test significance; n.s., not significant; *\*\*\*\*P <* 0.0001. (**C**) Confocal MIPs of p53-HaloTag-V5 knockin HEK293 cells transduced with Streptavidin-mtagBFP2-NES lentiviral particles and then incubated with 100nM JF_635-_HTL (steady-state) or 100nM dtb-JF_635-_HTL (trapped) for 24 hours. For the release condition, cells labeled with 100nM dtb-JF_635-_HTL were then incubated with 80 µM biotin for 3 hours. Cells were fixed with 4% PFA, immunostained with rabbit anti V5 and anti rabbit Alexa Fluor 555 secondary antibody, and stained with SYTO16 DNA dye.

We next asked if BOLT is generalizable enough to manipulate the localization of an endogenous transcription factor. To test this, we utilized a HEK293 cell line containing a HaloTag-V5-tag knockin at the endogenous locus for P53 (heterozygous tagging). Cells expressing p53-HaloTag-V5-Tag were transduced with Streptavidin-mtagBFP2-NES lentiviral particles and then incubated with JF_635-_HTL or dtb-JF_635-_HTL overnight to test trapping. Confocal fluorescence microscopy revealed endogenous p53-HaloTag could indeed become trapped in the cytosol after incubation with dtb-JF_635-_HTL and could be released with 80 µM biotin incubation to restore nuclear localization (Fig. 5C). These experiments demonstrated BOLT can be used to trap nuclear proteins in the cytosol and to subsequently induce their nuclear import.

## Discussion

Controlling protein localization through chemically induced dimerization or induced proximity has been instrumental for mechanistic dissection and acute functional perturbation. Both established and more recently developed platforms, including FRB-FKBP^4–7^, SBP-streptavidin^12,14,15^, SAMBA^16^, and CATCHFIRE^18^ have enabled a broad range of applications, but differ in their inducibility, reversibility, labeling capabilities, and potential for baseline perturbation. Building on these existing concepts, BOLT integrates the broad utility of a genetically encoded HaloTag with multifunctional DTB-JF-HTLs and a streptavidin-based trap. This design allows a POI to retain its native localization and function before manipulation, undergo rapid trapping following addition of a DTB-JF-HTL, and subsequently be released by biotin competition, restoring the POI’s original localization and function. BOLT may therefore be particularly advantageous for endogenous HaloTag knock-in proteins, for which minimizing perturbation before the experimental intervention is essential. More broadly, BOLT expands the available modalities for acute, reversible protein and organelle manipulation in cells while enabling high-precision visualization using a genetically encoded HaloTag, a streptavidin-based trap, and two bioavailable small molecules, DTB-JF-HTL, and biotin.

One of BOLT’s central features is that the multifunctional HTLs serve simultaneously as inducible dimerization reagents and as bright, persistent fluorescent labels. HaloTag fusion proteins can be pulse-labeled before trapping and followed continuously during both redistribution and biotin-mediated release. Importantly, the nature of covalent association of HTL and the HaloTag^38^ retains DTB-JF-HTL with POI after release from the streptavidin trap, enabling direct measurement of redistribution kinetics, recovery of steady-state localization, and the fate of the manipulated protein population. When combined with the growing repertoire of HaloTag ligands, BOLT can also be adapted to label and manipulate distinct POI cohorts. As demonstrated with MAN2A1-HaloTag knock-in RPE1 cells, sequential pulse-chase labeling with distinct JF-HTLs can distinguish temporally defined protein cohorts. This feature creates opportunities to compare newly synthesized and pre-existing POI populations; determine whether protein age influences trafficking, retention, or degradation; and resolve whether different cohorts respond differently to an acute relocalization perturbation. The development of additional spectrally separable DTB-JF-HTLs would further enable multiplexed cohort tracking and could reveal differences between newly synthesized and older proteins, or between organelles of different ages or metabolic histories.

Our experiments also illustrate the potential of BOLT to reveal previously inaccessible aspects of protein trafficking. Using endogenous MAN2A1-HaloTag, we observed a gradual Golgi-to-ER redistribution of a full-length Golgi glycosylation enzyme and this ER-retrieved population remained Golgi-destined once the trapping was reversed, supporting the hypothesis that some enzymes constitutively cycle between Golgi and ER. These findings raise several questions worthy of further interrogation: What is the physical nature of the transport vessel for Golgi-to-ER recycling? How is this recycling regulated and how are cargos selected for recycling? What is the biological significance of this pathway? Answering these questions may provide insights into the quality control of Golgi enzymes and how Golgi remodels during neuronal activity^39,40^, cell migration^41^, and chemotherapy in colorectal cancer cells^42,43^.

The observed mitochondrial fragmentation during BOLT-induced minus-end-directed microtubule transport (Fig. 4B) was unexpected. One possible explanation is that sustained motor-driven pulling increased mitochondrial membrane tension, which would favor fission – however, the relationship between microtubule association, mechanical force, mitochondrial motility, and mitochondrial morphology appears to be context dependent^44–46^. Going forward, it will be important to determine whether trafficking-associated fragmentation coincides with increased ER-mitochondria contact sites and local enrichment of dynamin-related GTPase DRP1 because these factors also play a role in mitochondrial fission^47^. BOLT also revealed differential hitchhiking behavior between peroxisomes, LDs, and mitochondria (Fig. 4F–H, Extended Data 5A, 5B). While peroxisomes traveled along with mitochondria moving along the microtubule network, LDs remained stationary during this process. This differential hitchhiking behavior likely reflects the strength of the organelles’ interaction: while peroxisomes are stably tethered to mitochondria, LDs may form more dynamic interaction with mitochondria.

There are some limitations and considerations of the current implementation that one should keep in mind. First, because streptavidin is tetrameric, membrane-tethered streptavidin can promote unintended higher-order clustering^48^. In our hands, membrane-anchored streptavidin alone often induced aggregation of organelles such as the ER and mitochondria, except at very low expression levels (data not shown). This may limit the use of the current trap configuration for experiments involving organelle–organelle interactions. Monomeric streptavidin may help mitigate this limitation, although its lower affinity would likely require further optimization before use^49^. Second, DTB-JF-HTLs can associate directly with streptavidin in the absence of a HaloTag. Although DTB-JF_635-_HTL exhibits increased fluorogenicity upon HaloTag binding, DTB-JF_646-_HTL remains fluorescent when bound to streptavidin alone. This background signal can complicate quantitative interpretation of the distribution or abundance of a HaloTag-fused POI. We addressed this concern using two complementary strategies: (1) pre-labeling HaloTag-fused POIs, washing away unbound ligand, and subsequently inducing the streptavidin trap; and (2) independently monitoring the POI with a tandem fluorescent protein tag or immunofluorescence. Future development of additional fluorogenic DTB-JF ligands should improve the quantitative performance of BOLT. Finally, although the basal concentration of biotin in media such as DMEM/F12 (14.3 nM) did not cause noticeable interference/leakiness of the BOLT system in RPE1 or HEK293 cells, other media formulations or animal tissues may contain higher biotin concentrations, potentially requiring biotin-depleted media for efficient trapping to occur. Beyond the development of spectrally distinct DTB-JF-HTLs, an important next step for further BOLT optimization will be the creation of orthogonal multifunctional HTL systems that use independent trapping and release mechanisms. Such systems could enable logical control of perturbations, including sequential, combinatorial, or conditionally gated manipulation of distinct proteins, organelles, or protein cohorts within the same cell. Recently developed *de novo* designer JF dye binders, such as NovoTag^50^, may provide an alternative to HaloTags that could further simplify implementation of BOLT, particularly for endogenous genome engineering due to their smaller size.

Despite these considerations, we believe BOLT will be widely applicable to exogenous HaloTag-compatible systems, allowing researchers to support existing and establish new generalizable mechanistic principles in cell biology. Looking forward, we envision that BOLT’s greatest utility lie in endogenous settings. Technically, BOLT is easier to implement as single HaloTag knock-in can support both visualization and reversible trap-and-release, whereas several alternative approaches require separate tagging modules for manipulation and imaging. Functionally, BOLT has no trapping activity at steady state minimizing perturbation of the POI before DTB-JF-HTL addition. This property is important for endogenous proteins, whose localization, complex assembly, and/or activity may be sensitive to chronic recruitment or constitutive dimerization.

Because BOLT can be introduced into any endogenous HaloTag model through exogenous expression of a streptavidin trap in principle (e.g., HaloTag knock-in induced pluripotent stem cell (iPSC)-related resources – such as cells from the iSPC Neurodegenerative Disease Initiative), it may allow investigators to repurpose existing HaloTag-tagged alleles without further genomic engineering^51^.

We have shown BOLT is a versatile, integrated platform that will allow us to move beyond descriptive localization studies and continue pursuing experimentally controlled tests of how protein and organelle distribution shapes cellular function. By coupling rapid, reversible trapping with persistent fluorescent labeling, BOLT enables perturbation of a defined protein or organelle population while directly tracking its behavior during both displacement and recovery. More broadly, BOLT provides a generalizable framework for converting spatial correlations into experimentally testable causal relationships between protein and organelle organization and cellular function.

## Acknowledgements

We would like to thank Danny M. D’Amore for scientific editing, Pratik Kumar for technical support for chemical synthesis, the Vector Laboratory and Protein Production Facility at St. Jude Children’s Research Hospital (SJCRH) for generating lentiviral particles and recombinant msGFP-HaloTag, respectively, Marquetta Nebo, Jennifer Maglisco, and Monique Payton for administrative and operational assistance.

## Author Contributions

W.C.B. and C-L.C. conceived the project; P.K. and S.P. generated the DTB-JF-HTLs and biotin-JF-HTLs; L.D.L. supervised DTB-JF-HTL design and production; W.C.B. generated plasmids; W.C.B., K.R.B., and M.R.H. performed image acquisition and analysis; W.C.B., J.L.S., and J.D.V. performed in-gel fluorescence assays; M.S.H. generated and validated the RPE1 knock-in cells, which was supervised by S.P-M.; H.K.K generated and validated the HEK293 knock-in cells, which was supervised by A.C.; W.C.B. and C-L.C. wrote and drafted the manuscript with input from all authors. C-L.C. supervised the project.

## Materials and Methods

### Chemical compounds

JF_635-_HTL was purchased from Tocris (88-081-0U). JF_549-_HTL, JF_646-_HTL, JFX_549-_HTL, JFX_646-_HTL, biotin-JF_635-_HTL and biotin-JF_646-_HTL were available from previous work^22,52,53^. DTB-JF_635-_HTL and DTB-JF_646-_HTL were synthesized by coupling 3”-carboxy-JF_635-_HTL or 3’-carboxy-JF_646-_HTL with DTB-PEG_2-_NH ^22,54,55^

### Production and titration of lentiviral vectors

Production and titration of lentiviral vectors were performed as described previously^56,57^. Briefly, SJ293TS-DPB cells were transfected with the transfer vector and the helper plasmids pCAG-kGP1-1R-AF, pCAG-VSVG-AF, and pCAG4-RTR2-AF using PEIpro (Polyplus Transfection) and grown in Freestyle 293 Expression media (Thermo Fisher Scientific) supplemented with 20% LV-MAX™ media (Thermo Fisher Scientific) at 37°C with 8% CO_2 a_nd shaking at 125 rpm. Viral vector–containing supernatants were collected 48 h post-transfection, clarified by centrifugation at 330 × *g* for 5 min, and filtered (0.22-µm filter). Vector preparations were concentrated using an Amicon® Ultra Centrifugal Filter with a 100-kDa molecular weight cutoff (Millipore Sigma) as per the manufacturer’s instructions to achieve a 10-fold reduction in volume. Vector preparations were centrifuged at 3,000 × *g* for approximately 5 min. The concentrated vector preparation was 0.22-µm sterile filtered, aliquoted, and stored at −80°C until use.

Lentiviral vector titration was performed by transducing HOS-6A5 cells (derived from ATCC CRL-1543) with serially diluted vector preparations in the presence polybrene (5–8 µg/mL; Millipore Sigma), and were grown in DMEM (Corning) supplemented with 10% fetal bovine serum (FBS; Seradigm) and 2 mM L-alanyl-L-glutamine (Corning) at 37°C with 5% CO_2._. Four days post-transduction, genomic DNA was isolated from transduced cells and after digestion with the restriction enzyme *Msp*I, was used as a template in PCR using a QX200 Droplet Digital PCR System (Bio-Rad). The following primer-probe sets were used to amplify the HIV *psi* sequence and the endogenous control gene, *RPP30*, 5’-ACTTGAAAGCGAAAGGGAAAC-3’, 5’-CACCCATCTCTCTCCTTCTAGCC-3’ and probe 5’-56-FAM-AGCTCTCTC-ZEN-GACGCAGGACTCGGC-3IABkFQ-3’ and 5’-GCGGCTGTCTCCACAAGT-3’, 5’-GATTTGGACCTGCGAGCG-3’ and probe 5’-5HEX-CTGACCTGA-ZEN-AGGCTCT-3IABkFQ-3’, respectively. Vector titers and copy numbers were determined by calculating the number of copies of HIV *psi* to every two copies of *RPP30*, multiplied by the number of cells transduced, and if necessary, multiplied by the dilution factor.

### Plasmids and cloning

pEF_ef1a_cGFP-HaloTag was generated for the permeability assay via PCR amplification of the cGFP-HaloTag gene block and In-Fusion cloning into a pEF-GFP backbone that was linearized with *Kpn*I (Thermo Scientific, ER0521) and *Not*I (Thermo Scientific, FERFD0594). All In-Fusion cloning was performed by linearizing the vector with restriction enzymes or by PCR and PCR amplifying the insert to have 15–20 base pairs of overlapping sequence with the vector at the 5’ and 3’ ends and performing the In-Fusion reaction with the In-Fusion enzyme snap assembly master mix (Takara, 638948) according to the manufacturer’s instructions. pET SUMO-msGFP-HaloTag was generated for recombinant msGFP-HaloTag expression by In-Fusion cloning *Escherichia coli* codon-optimized msGFP-HaloTag gene block into pET_His6-Sumo-LIC (2S-T) backbone linearized with *Ssp*I (Thermo Scientific, ER0771).

To generate SJL12 CMV streptavidin-KDEL IRES TNFα-SBP-HaloTag (used as a cloning intermediate in this study), TNFα-SBP-HaloTag was amplified from Halo-TNFa-RUSH (Addgene,166901) to generate *Age*I and *Bsr*GI ends, digested with *Age*I and *Bsr*GI (Thermo Scientific, FD0934) and ligated into SJL12 CMV-GFP (which was also digested with *Age*I and *Bsr*GI^58^). SJL12 CMV streptavidin-KDEL IRES TNFα-HA-mNG-HaloTag was generated by linearizing SJL12 streptavidin-KDEL IRES TNFα-SBP-HaloTag with mutagenic primers to replace SBP with an HA tag, and an mNG insert generated by PCR with homologous 5’ and 3’ ends was added via In-Fusion cloning. Man2-N’ refers to a synthetic minimal Golgi marker derived from the mouse MAN2A1 gene, containing N terminal residues which are sufficient to localize to Golgi. The SJL12 streptavidin-KDEL IRES Man2-N’-SBP-mCherry RUSH vector was generated by digesting the SJL12 backbone and Str-KDEL_ManII-SBP-mCherry (Addgene, 65253) with *Mlu*I and *Pac*I followed by T4 ligation^12^. SJL12 CMV Streptavidin-KDEL IRES Man2-N’-HA-mNG-HaloTag was generated by cutting SJL12 CMV Streptavidin-KDEL IRES TNFα-HA-mNG-HaloTag with AscI and BsrGI to remove TNFα-HA-mNeonGreen and replace it with a gene fragment encoding Man2-N’ (1-102)-HA-mNeonGreen using the same enzymes and T4 ligation. SJL12 streptavidin-KDEL was generated by using *Not*I restriction digestion to remove the IRES TNFα-SBP-HaloTag from SJL12 streptavidin-KDEL IRES TNFα-SBP-HaloTag and T4 ligation to close the vector. Doxycycline inducible SJL21i streptavidin-KDEL EF1α PuroR-P2A-Tet-On 3G was generated by linearizing SJL21i TRE-Cherry EF1α PuroR-P2A-Tet-On 3G (provided by the St. Jude Children’s Research Hospital [SJCRH] Vector Laboratory) with *Asc*I (Thermo Scientific, FD1894) and *Pac*I (Thermo Scientific, FD2204) and inserting a streptavidin-KDEL PCR fragment using In-Fusion.

SJL12 streptavidin-mtagBFP2-Sec61 was generated by GenScript’s cloning service through restriction digestion of SJL12 CMV-GFP with *Eco*R1 (Thermo Scientific, FD0274) and *Not*I and ligating a synthetic gene fragment encoding streptavidin-mtagBFP2-Sec61. SJL12 streptavidin-mtagBFP2-NES was generated by mutagenesis PCR with primers designed to exclude Sec61 and add NES from the HIV Rev protein (copied from the PiggyBac-APEX-V5-NES plasmid map; Addgene, 462274), and the linear PCR was closed by In-Fusion^59^. SJL12 flag-BicD2-594-streptavidin was generated using GenScript’s FLASH gene service. A synthetic gene fragment was designed containing flag-BicD2-594 copied from pBa-eGFP-flag-BicD2-594-FKBP (Addgene, 63569), a 3X SAGG linker, and streptavidin, which was inserted into SJL12 CMV-GFP digested with *Age*I (Thermo Scientific, ER1461) and *Not*I^32^. pGenlenti TOM20-HaloTag IRES PuroR was generated by GenScript^60^. The PMP34-HaloTag mammalian expression plasmid was described previously^37^. Doxycycline-inducible SJL21i TRE HaloTag-ATF4 EF1α PuroR-P2A-Tet-On 3G was generated by digesting SJL21i TRE-Cherry EF1α PuroR-P2A-Tet-On 3G with *Asc*I (Thermo Scientific, FD1894) and *Pac*I (Thermo Scientific, FD2204) and inserting two PCR fragments, HaloTag (from TOM20-HaloTag) and *ATF4* (amplified from human complementary DNA) using In-Fusion cloning. SJL12 CMV mCherry-SKL was generated by first subcloning an mOrange2-SKL insert from mOrange2-SKL expression plasmid into SJL12 CMV GFP using *Age*I and *Not*I restriction sites, and then swapping mOrange2 for mCherry by cutting and pasting an mCherry insert from mCherry-C1 using *Age*I and *Bsr*G1 restriction sites.

PCRs for amplifying inserts or linearizing plasmids were performed using Phusion Hot Start 2 DNA polymerase (Thermo Scientific, F549S) according to the manufacturer’s instructions. Restriction digestions were performed by digesting 1 µg of plasmid with 1 µL (10 enzyme units) of Thermo Scientific restriction enzymes. Digested plasmids and PCR products were run in 1% agarose (Thermo Scientific, A9369) gels with 1X TAE buffer (Invitrogen, 24710030) and SYBR Safe DNA dye (Invitrogen, S33102). Bands were excised and DNA was purified using the NucleoSpin Gel and PCR Clean-up kit (Macherey-Nagel, 740609.250) per the manufacturer’s instructions. T4 ligations were performed with purified DNA using T4 DNA ligase (Thermo Scientific, EL0011) according to the manufacturer’s instructions. After ligation or infusion reactions, the plasmids were transformed into Stellar competent *E. coli* cells (Takara, 636766), plated on Luria-Bertani (LB) plates containing 40 µg/mL kanamycin (MOLTOX, 50948843) or 50 µg/mL carbenicillin (Sigma-Aldrich, C1389-5G) for selection. Plasmids were purified using the NucleoSpin Plasmid Transfection-grade Mini Kit (Macherey-Nagel, 740490.250) for transfection plasmids or NucleoBond Xtra Midi EF Kit (Macherey-Nagel, 740420.50) for lentiviral production plasmids, both per the manufacturer’s instructions. All vectors were confirmed by whole plasmid sequencing. Plasmids are available upon request. Primer sequences and cloning schemes are listed in Table S1.

### Cell culture and transduction

hTERT RPE-1 cells (referred throughout as RPE1) were originally sourced from ATCC (CRL-4000). Their identity was authenticated using short tandem repeat (STR) profiling by the Center for Advanced Genome Engineering and routinely tested for mycoplasma using MycoAlert detection kits (Lonza, LT07-318). RPE1 cells were maintained in DMEM/F12 media (Gibco, 30-2006) with 10% FBS (Gibco, A5669801), 1% penicillin/streptomycin (Gibco, 15140122), and 0.01 mg/mL hygromycin B (Invitrogen, 10687010), grown in T25 flasks and split 1:10 every 3 days. HEK293 cells were maintained in DMEM (Gibco, 11995073) with 10% FBS and 1% penicillin/streptomycin. p53-HaloTag-V5-pt2a-NrsR knockin HEK293 cells were maintained in DMEM with 10% FBS, 1% penicillin/streptomycin, 1% glutamax, and 100 µg/mL Nourseothricin (N-500-100, GOLDBIO) selection for maintentance of the HaloTag knockin.

For doxycycline-inducible lentiviral vectors containing puromycin resistance (SJL21i Streptavidin-KDEL EF1α PuroR-P2A-Tet-On 3G and SJL21i HaloTag-ATF4 EF1α PuroR-P2A-Tet-On 3G), transduced cells were maintained with 6 µg/mL puromycin (Gibco, A1113803) for stable expression. For transductions, 4000 cells were seeded per well in 8-well chambers (Lab-Tek Z734853-96-EA) with 300 µL media per well. After 24 h, the media was replaced with fresh media containing 10 µg/mL protamine sulfate (MP Biomedicals 194729) and a corresponding volume of concentrated lentivirus to achieve 20–150 MOI (MOIs were individually optimized for each lentivirus). Cells were imaged by spinning disc confocal microscopy as described below 48–72 h after transduction after replacing the media.

### Generation of MAN2A1-HaloTag/LMAN-mEmerald RPE1 knock-in cell line

MAN2A1-HaloTag/LMAN1-mEmerald RPE-1 cells were made using CRISPR technology in the CAGE (RRID:SCR_027027). In brief, genome edits were introduced sequentially. The LMAN1-N-terminal mEmerald knock-in was generated first, and a clonal cell line was isolated and validated before introduction of the MAN2A1-HaloTag knock-in. For each editing step, 200,000–400,000 RPE-1 cells (provided by the CAGE) were transiently co-transfected with pre-complexed ribonucleoprotein consisting of 100 pmol chemically modified sgRNA (IDT) and 50 pmol 3×NLS SpCas9 protein (SJCRH Protein Production Facility), together with 500 ng dsDNA donor template. Transfection was performed via nucleofection (Lonza, 4D-Nucleofector™ X-unit) using solution P3 and program EA-104 in a small (20-µL) cuvette according to the manufacturer’s protocol, and cells were plated in prewarmed (37°C) media. Five days post nucleofection, cells were single-cell sorted by flow cytometry (SJCRH Flow Cytometry and Cell Sorting Shared Resource) on mEmerald (LMAN1) or JF_646-_HTL (MAN2A1) into 96-well tissue culture treated plates containing pre-warmed media. Cells were clonally expanded and screened for the desired modification via PCR-based assays and targeted amplicon sequencing via next generation sequencing as previously described^61^. Correctly edited clones were expanded and sequence confirmed. The LMAN1 N-terminal mEmerald knock-in clone used for subsequent MAN2A1-HaloTag targeting contained one correctly tagged *LMAN1* allele and one *LMAN1* knockout allele. For *MAN2A1*, only correctly tagged alleles were identified following HaloTag integration in the final double-tagged clone. Final clone cell identity was authenticated using the PowerPlex® Fusion System (Promega) performed at the Hartwell Center (SJCRH). All clones tested negative for mycoplasma using the MycoAlertTMPlus Mycoplasma Detection Kit (Lonza, LT07-710). Genome editing reagents and sequencing primers listed in Table S2.

Editing construct sequences and screening primers are listed in Table S2. Deep sequencing adaptors and knock-in modification are in upper case.

### Generation of p53-HaloTag-V5-PT2A-3xNsrR HEK293 knock-in cell line

*The p53-HaloTag-V5-PT2A-3xNsrR* (shortened to p53-HaloTag-V5 in the results) knock-in cell line was generated following a previously reported method with minor modifications^62,63^. Briefly, HEK293T cells were seeded in 24-well plates and transfected the following day with 100 ng pCAS (Addgene plasmid #206994), 50 ng pDNR-gRNA (Addgene plasmid #206990), the 50 ng target gRNA plasmid, and the corresponding 500 ng pDNR-GGGS_HaloTag_V5_PT2A_3xNrsR donor plasmid (Addgene plasmid #263453, 263454, or 263456). Transfections were performed using Lipofectamine 2000 (Invitrogen, 11668027) according to the manufacturer’s instructions. The media was replaced the following day and cells were split upon reaching confluence every 2-3 days. On day 7 after transfection, cells were transferred to media containing 100 μg/mL nourseothricin (N-500-100, GOLDBIO) for selection and continued to be spilt in a similar fashion as before. Selection was continued until the parental, non-tagged cells were completely eliminated, which typically required approximately 2 weeks. Proper junction formation was confirmed by PCR using genomic DNA extracted with QuickExtract DNA Extraction Solution (LGC Biosearch Technologies, LGCQE09050). Primers were designed to amplify the junction between the endogenous p53 locus and the integrated HaloTag donor sequence. The expected PCR products were analyzed by agarose gel electrophoresis, and correct junction formation was further confirmed by Sanger sequencing (Genewiz). gRNA sequence and primers used to amplify and sequence the TP53-HaloTag junction are listed in Table S3.

### Spinning disc confocal microscopy

All microscopy images were captured using a Nikon Ti2 microscope with Yokogawa CSU-W1 Spinning Disk unit, XY galvo scanning module, and Tokai Hit STXG CO_2 i_ncubation system using 60x (CFI APO TIRF, NA 1.49 oil) or 40x (CFI60 PLAN APOCHROMAT LAMBDA D, NA 0.95 air) objectives.Stage incubation was set to 37°C and 5% CO2 for live cell imaging. For z-stacks, a 0.3-µm interval was used and 11–15 slices were taken. Z-stacks were processed into maximum z projections in Fiji ImageJ 2.18.0/1.54p^64^.

### HTL permeability assays

HEK293 cells were seeded in 12-well plates (1:10 split from a confluent 10-cm dish). Twenty-four hours later, each well was transfected with 25 ng pEF_msGFP-HaloTag (cytosolic) by mixing 12 µL Lipofectamine LTX Reagent with PLUS (Thermo Fisher, 15338100) diluted in 300 µL Opti-MEM (Gibco 31985062)with 6 µL PLUS reagent with 25 ng plasmid diluted in 300 µL Opti-MEM for a total volume of 600 µL. Twenty-four hours after transfection, cells were pulsed with the corresponding JF-HTL. After 2 h, 1 µM JF549-HTL was added for 30 min to react to any leftover HaloTag. Cells were harvested with 150 µL RIPA buffer (Santa Cruz; SC-24948) and 55 µL Laemmli/BME buffer (Bio-Rad; 1610747 and 1610710). Samples were heated at 50°C for 5 min before a 30-µL aliquot of sample was electrophoresed in tris glycine gels (Bio-Rad, 5671034) and imaged with aChemiDoc MP (BioRad, 12003154) per manufacturer’s instructions to measure in-gel fluorescence.

### DTB-JF_635-_HTL saturation curve in MAN2A1-HaloTag knock-in RPE1 cells

MAN2A1-HaloTag knock-in RPE1 cells were seeded at 10,000 cells per well in 8-well Lab-Tek chambers (Nunc, Z734853) with 300 µL media. The next day, the media was replaced with media containing DTB-JF_635-_HTL at concentrations of 0, 1, 10, 100, 250, 500, or 1000 nM. Twenty-four hours later, cells were fixed with 4% PFA (Electron Microscopy Services, 15710) in PBS for 15 min, washed twice with 300 µL PBS, and then stained with 1 µM JF549-HTL for 1 h at room temperature. Wells were washed twice with 300 µL PBS (Gibco, 10010049) and imaged with spinning disk confocal microscopy (described above). DTB-JF_635-_HTL mean fluorescence intensity was measured in Fiji version 2.18.0/1.54p with background subtraction. Finally, HaloTag saturation was quantified in GraphPad Prism version 11.1.0 and a nonlinear regression curve fitting was applied using the “binding – saturation one site total” option to model a saturation curve.

### Recombinant msGFP-HaloTag protein expression

The target protein was expressed in *E. coli* BL21(DE3). Cultures were grown in LB medium supplemented with ampicillin (100 µg/mL) at 37°C until an optical density OD_600 o_f 0.5–0.6 was reached. Protein expression was induced by the addition of 0.2 mM isopropyl β-D-1-thiogalactopyranoside.Ccultures were then incubated overnight 25°C. Cells were harvested by centrifugation at 5,000 rpm for 15 min at 4°C and the resulting cell pellets were stored at −20°C until further use.

### Affinity purification and tag cleavage

The cell pellet was resuspended in lysis buffer consisting of 25 mM Tris (pH 7.2), 200 mM NaCl, and 1 mM tris(2-carboxyethyl)phosphine (TCEP), supplemented with cOmplete™ protease inhibitor cocktail (Roche, 11873580001) and a *Serratia marcescens* NucA nuclease produed in-house. Cells were lysed via high-pressure homogenization in a PandaPlus homogenizer (GEA), and the lysate was clarified by high-speed centrifugation at 16,000 rpm for 60 min at 4°C.

The supernatant was loaded onto a nickel-affinity column equilibrated with the lysis buffer. The column was washed thoroughly with wash buffer (25 mM Tris (pH 7.2), 200 mM NaCl, 1 mM TCEP, and 25 mM imidazole) to remove non-specifically bound proteins. Target proteins were subsequently eluted using an elution buffer containing 25 mM (Tris pH 7.2), 200 mM NaCl, 1 mM TCEP, and 250 mM imidazole.

To remove the affinity tag, the pooled elution fractions were digested overnight at 4°C by adding TEV protease (produced in-house).

### Size exclusion chromatography

Following tag cleavage, the sample was concentrated and further purified by size exclusion chromatography on a Superdex S-200 26/60 column (Cytiva). The column was equilibrated and eluted with a buffer comprising 25 mM Tris (pH 7.2), 200 mM NaCl, and 1 mM TCEP. Fractions corresponding to the monodisperse peak of the target protein were collected, pooled, and verified for purity via SDS-PAGE.

### Recombinant HaloTag pulldown assays

Pulldowns were performed with 1 µg of recombinant msGFP-HaloTag diluted in PBS with 0.2% TX100 (Sigma, 93443)for a total reaction volume of 100 µL. First, msGFP-HaloTag was reacted with the corresponding JF-HTL at a concentration of 1 µM for 30 min at room temperature before being stored at 4°C overnight. Pierce Streptavidin Magnetic Beads (Thermo Scientific, 88817) were washed twice with PBS 0.2% TX100 at room temperature to remove storage buffer (1 mL/500 µL bead slurry). Before starting the pulldown, excess sample was used as input.

Then, 30 µL bead slurry was added to 1 µg msGFP-HaloTag-JF-HTL for a working volume of 130 µL and rotated end over end for 2 h at 4°C. After collecting the flow through supernatant, beads were washed three times with 250 µL PBS 0.2% TX100. Beads were then resuspended in 100 µL of 160 µM biotin (Invitrogen, B1595) in PBS 0.2% TX100 for the first elution for 20 min at room temperature. After collecting the supernatant (biotin elution), beads were washed twice with 250 µL PBS 0.2% TX100 and then resuspended in 100 µL SDS Laemmli sample buffer (BioRad, 1610737) diluted 1:2 with PBS 0.2% TX100 to 1X concentration (∼1% SDS) with 1:20 2-Mercaptoethanol (BioRad, 1610710) and incubated at 60°C for 10 min for the second elution. Input, flowthrough, biotin elution, and sample buffer elution were diluted 1:2 in 2X sample buffer, boiled for 5 min at 95°C, and electrophoresed on Tris glycine 4%–20% gels (BioRad, 4561096), and JF-HTL in-gel fluorescence was detected using a ChemiDoc MP (BioRad, 12003154) imager.

### Comparing HTL binding ability to Streptavidin in RPE1 cells

Approximately 4000 RPE1 cells were seeded per well in 8-well Lab-Tek imaging chambers with 300 µL media per well as described in the Cell culture and transduction section. The next day, they were transduced with SJL12 streptavidin-mtagBFP2-NES lentivirus encoding cytosolic streptavidin. Forty-eight hours after transduction, the media was replaced with 270 µL fresh media containing 100 nM HTL and cells were allowed to incubate for 1 h to allow the HTL to permeate the cells and bind the streptavidin. Then, 30 µL of 800 µM biotin diluted in media (or just media as a control) was added to each corresponding well for an 80-µM final concentration to test releasability of each HTL for 30 min. Cells were then washed with PBS, fixed with 4% PFA in PBS for 15 min, washed twice with 300 µL PBS, and imaged with spinning disk confocal microscopy (described above). Cells were manually encircled using the freehand selection tool and added to the ROI manager in Fiji ImageJ 2.18.0/1.54p. Mean fluorescence intensity was measured per cell with background subtraction by subtracting minimum fluorescence intensity per cell.

### Secretory cargo trafficking BOLT experiments

RPE1 cells were seeded 4000 cells/well in Lab-Tek 8-well chambers with 300 µL media. After 24 hours, cells were transduced with a secretory cargo lentiviral vector, SJL12 streptavidin IRES Man2-Nter-mNG-HaloTag or SJL12 streptavidin IRES TNFα-HA-mNG-HaloTag. For the BOLT versus RUSH comparison, SJL12 streptavidin IRES Man2-Nter-SBP-mCherry was co-transduced simultaneously with SJL12 streptavidin IRES Man2-Nter-mNG-HaloTag. For endogenous MAN2A1-HaloTag experiments, only the trap, SJL12 streptavidin-KDEL, was transduced. Forty-eight hours after transduction, the media was replaced with media containing 100 nM JF_635-_HTL (Tocris, 8808) (steady state condition) or 100nM DTB-JF_635-_HTL (trapped condition) and cells were incubated overnight to label a population of secretory cargo. On the day of imaging, media was replaced with DMEM/F12 10% FBS 1% penicillin/streptomycin media lacking phenol red and the chambers were imaged with spinning disc confocal microscopy with stage incubation set to 37°C and 5% CO_2._ For biotin release conditions, an 800-µM biotin 10X stock was made up in media, and 30 µL was added to each well for an 80-µM final concentration. Z stacks were acquired for each condition, and maximum intensity projections were made for analysis. For biotin release timelapses, biotin was added immediately after the first frame of 2D acquisition. For endogenous MAN2A1-HaloTag experiments, cells were fixed with 4% PFA for 15 min at room temperature, washed twice with PBS, permeabilized with 0.2% TX100, 1% BSA, and 5% goat serum in PBS (staining buffer) and immunostained with mouse anti-Golgin97 (eBioscience, 14-9767-82) for 1 h. Then, cells were washed twice with PBS and stained with secondary goat anti-mouse Alexa Fluor 488 (Invitrogen, A-11029), before washing twice more with PBS and imaging. To analyze the max-to-mean intensity ratio, cells were manually encircled using freehand selection in Fiji, added to the ROI manager, and mean, minimum, and maximum intensities per cell were measured. Max/Mean was calculated as follows: (max intensity – min intensity) / (mean intensity – min intensity). Data was plotted using GraphPad Prism version 11.1.0.

### MAN2A1-HaloTag recycling experiments

MAN2A1-HaloTag LMAN1-mEmerald dual knock-in RPE1 cells were transduced with a doxycycline-inducible lentiviral vector, SJL21i TRE streptavidin-KDEL Ef1a Tet3ON-T2A-PuroR (100 MOI) and maintained in media containing 6 µg/mL puromycin. For each experiment, 4000–8000 cells were seeded per well of 8-well chambers for 48 h with normal media lacking puromycin to minimize unnecessary stress. The night before imaging, media was replaced with fresh phenol red free media containing 100 nM JFX_646-_HTL (negative control) or 100 nM DTB-JF_646-_HTL to pulse MAN2A1-HaloTag for 18 h. The next morning, the HTLs were aspirated and any unreacted HaloTag was quenched with 10 µM 7-bromo-1-heptanol (Thermo Scientific Chemicals, H54762.03) diluted in 200 µL media in the cell culture incubator set to 37°C and 5% CO2 for 10 min^65^. Then, each well was washed three times (once briefly and twice for 5 min each) with 300 µL media in the cell culture incubator to thoroughly remove excess HTL or 7-bromo-heptanol. Then, media was replaced with media containing 100 nM JFX549-HTL to label newly synthesized MAN2A1-HaloTag ±1 µg/mL doxycycline to induce expression of streptavidin-KDEL to trap the recycling population of MAN2A1-HaloTag-DTB-JF_646-_HTL for a 6-h chase. Cells were imaged 6 h after doxycycline induction to measure efficiency of MAN2A1-HaloTag-DTB-JF_646-_HTL trapping in the ER. Then, timelapses were recorded to assess the molecule’s capacity to traffic from the ER back to the Golgi after the addition of biotin (for a final concentration of 80 µM). Images were quantified by measuring the max-to-mean intensity ratio of individual cell ROIs in Fiji ImageJ 2.18.0/1.54p.

### BICD2-streptavidin–mediated organelle trafficking

For mitochondria trapping experiments, wild-type hTERT-RPE1 cells were seeded in 8 well Lab-Tek chambers at 4000 cells per well with 300 µL of media. The next day, each well was transduced with the lentiviral vectors pGenLenti TOM20-HaloTag IRES PuroR (25 MOI) and SJL12 flag-BicD2-streptavidin (50 MOI), and for peroxisome imaging experiments, SJL12 mCherry-SKL (10–20 MOI). Forty-eight hours later, wells were stained with 100 nM Mitotracker Green FM (Invitrogen, M7514) and 100 nM monodansylpentane (i.e., AUTODOT™ Visualization Dye, SM1000b; BTL BioTechno Labs) in Fluorobright DMEM (Gibco, A1896701) containing 5% FBS for 5–10 min, aspirated, and then media was replaced with fresh Fluorobright media containing 5% FBS with the corresponding HTL at a working concentration of 100 nM. Cells were then imaged using spinning disc confocal microscopy with 60x oil immersion lens. For statistical quantification, z stacks were acquired and processed to maximum z projections in Fiji ImageJ 2.18.0/1.54p, and max-to-mean intensity ratios were measured for manually encircled cell ROIs. mCherry-SKL images were pre-processed with the ‘subtract background’ function in Fiji using a rolling ball radius of 50.0 px to minimize cytosolic background signal. For the control, untrapped condition, JF_635-_HTL (Tocris, 88-081-0U) was incubated with cells overnight. For the nocodazole condition, cells were pretreated with 30 µM nocodazole (Sigma Aldrich, 487928) for 2 h to depolymerize microtubules before adding DTB-JF_635-_HTL for 30 min. For biotin reversal conditions, biotin was added to wells for a final concentration of 80 µM after 30 min of trapping with DTB-JF_635-_HTL. For 2D timelapses, DTB-JF-HTL (final concentration of 100 nM) was added to the well immediately after the first frame to recruit Flag-BicD2-Streptavidin and biotin (final concentration of 80 µM) was added after 30 min to test DTB-JF-HTL binding reversibility.

To assess mitochondria morphological changes, we used the Mitochondria Analyzer plugin to measure the form factor and branch junctions per mitochondria^66^. Individual cell ROIs were cropped, and the ‘edit>clear outside’ command was used to eliminate signal outside of a single cell. Next, images were converted to 8-bit. Then, the ‘2D Threshold’ function was selected within the plugin, block size was set to 1.250, C-value was set to 5, and “automatically perform analysis?” was checked to threshold and simultaneously analyze each image.

Radial distribution analysis was performed using the ‘Radial Profile Extended’ plugin (developed by Philippe Carl; https://github.com/phcarl/Radial-Profile-Extended) in Fiji ImageJ 2.18.0/1.54p. Cells in maximum z projections were manually encircled using the freehand selection tool to generate an ROI and the centroid coordinates of that ROI were measured. The ‘edit>clear outside’ command was used to remove fluorescence outside of the ROI. Then, the ‘Radial Profile Angle’ plugin was opened and the radius was set to 700 px, which was large enough to encircle entire cells, and the circle was centered on the cell centroid. Then, “calculate radial profile” was clicked and “list” was selected to copy the radial profile data to a Microsoft Excel spreadsheet, which was plotted with GraphPad Prism 11.1.0.

For the peroxisome trapping experiment, cells were transduced with SJL12 flag-BicD2-streptavidin and SJL12 mCherry-SKL before being transfected with the PMP34-HaloTag expression vector the following day with 0.75 µL Lipofectamine 3000 and 0.5 µL P3000 reagent (Invitrogen, L3000015) according to the manufacturer’s instructions, with 100 ng plasmid per well. The following day, the cells were stained with 100 nM Mitotracker Green FM for 10 min, and then the media was replaced with Fluorobright DMEM with 5% FBS. Cells were then incubated with 100 nM DTB-JF-_635-_HTL for 60-min. Trapping was assessed by spinning disk confocal microscopy and flouresence was assessed using the max-to-mean intensity ratio in Fiji ImageJ 2.18.0/1.54p.

### HaloTag-ATF4 and p53-HaloTag-V5 cytosol sequestration

RPE1 cells were first transduced with doxycycline-inducible SJL21i HaloTag-ATF4 EF1α Tet3On-T2A-PuroR lentivirus and maintained with 6 µg/mL puromycin selection for at least 5 days before passage in puromycin containing media for stable selection. Then, for each experiment, cells were seeded at 4000 cells per well in 8-well chambers in media lacking puromycin. The next day, cells were transduced with SJL12 streptavidin-mtagBFP2-NES lentivirus. Forty-eight hours later, the media was replaced with 500 nM DTB-JF_635-_HTL (trapping condition) or JF_635-_HTL (no trapping condition) ±1 µg/mL doxycycline for 5 h to induce exogenous HaloTag-ATF4 expression. For the biotin release condition, biotin was added to a final concentration of 80 µM from an 800-µM working stock after the first 2 h of doxycycline induction. Five hours after doxycycline induction, cells were fixed with 4% PFA in PBS for 15 min at room temperature and washed two additional times with 300 µL PBS. To assess JF_635-_HTL signal specificity, cells were then immunostained with an anti-ATF4 antibody as an independent ATF4 label. For immunostaining, fixed cells were blocked and permeabilized with 0.2% TX100 in PBS with staining buffer (1% BSA and 5% goat serum [Gibco, 16210072]) for 30 min. Then, wells were incubated with 200 µL 1:200 rabbit anti-ATF4 (Cell Signaling Technology, D4B8) for 1 h, stained with 500 nM SYTO16 DNA stain (Invitrogen, S7578) for 10 min in PBS, washed twice with 300 µL PBS, and then incubated with 1:500 goat anti-rabbit Alexafluor 555 secondary antibody (Invitrogen, A32732) in staining buffer for 1 h. Then, wells were washed twice with 300 µL PBS and imaged via spinning disk confocal microscopy. For quantification, ATF4 mean fluorescence intensity was quantified in the nucleus and in the whole cell for each cell, background was subtracted by subtracting the minimum fluorescence intensity, and the nucleus to whole cell ratio was quantified to assess nuclear enrichment using Fiji ImageJ 2.18.0/1.54p.

For the biotin release timelapse video, HaloTag-ATF4 was induced with 1 µg/mL doxycycline for 5 h with with DTB-JF_635-_HTL to trap it in the cytosol. While recording the timelapse, biotin was added (to a 80-µM final concentration) immediately after the first frame to assess the release of HaloTag-ATF4.

For the p53-HaloTag cytosol sequestration experiment, 4000 p53-HaloTag-V5-PT2A-NrsR knockin HEK293 cells were seeded per well in 8 well Lab-Tek imaging chambers with 300 µL of media. 24 hours later, they were transduced with SJL12 CMV Streptavidin-mtagBFP2-NES lentiviral particles. The following day, the media was replaced with media containing 100 nM JF_635-_HTL or 100 nM dtb-JF_635-_HTL for overnight labeling and trapping. The following day, a final concentration of 80µM biotin was added to the release well for 3 hours. Then, wells were fixed with 4% PFA, washed twice with PBS, and immunostained with Rabbit anti V5 (Cell Signaling Technology; 13202S) with anti Rabbit Alexa Fluor 555 antibody as an independent marker for the HaloTag, and stained with SYTO16 DNA dye prior to confocal fluorescence imaging. Z-stacks were processed into maximum intensity projections in Fiji ImageJ 2.18.0/1.54p.

### Statistical analysis

For statistical comparisons, three or four independent experiments were performed for each experiment. For microscopy experiments, at least 6 cells were measured per replicate, and a mean was generated for each replicate. In GraphPad Prism, the experimental means were tested for statistical significance to assess reproducibility. For experiments with two conditions, Welch’s or paired *t*-tests were performed, while experiments with more than two conditions utilized a one-way ANOVA was with *post hoc* Tukey pairwise comparison testing. For grouped analysis involving more than two conditions, a two-way ANOVA was utilized with *post hoc* pairwise comparisons. Graphs are depicted as “superplots” with large points indicating the mean of an independent experiment and small points indicating individual cell values from all experiments, with bars indicating the mean +/-error (either standard deviation or standard error of the mean) for all experiments^67^.

---

**Extended Data 1.**
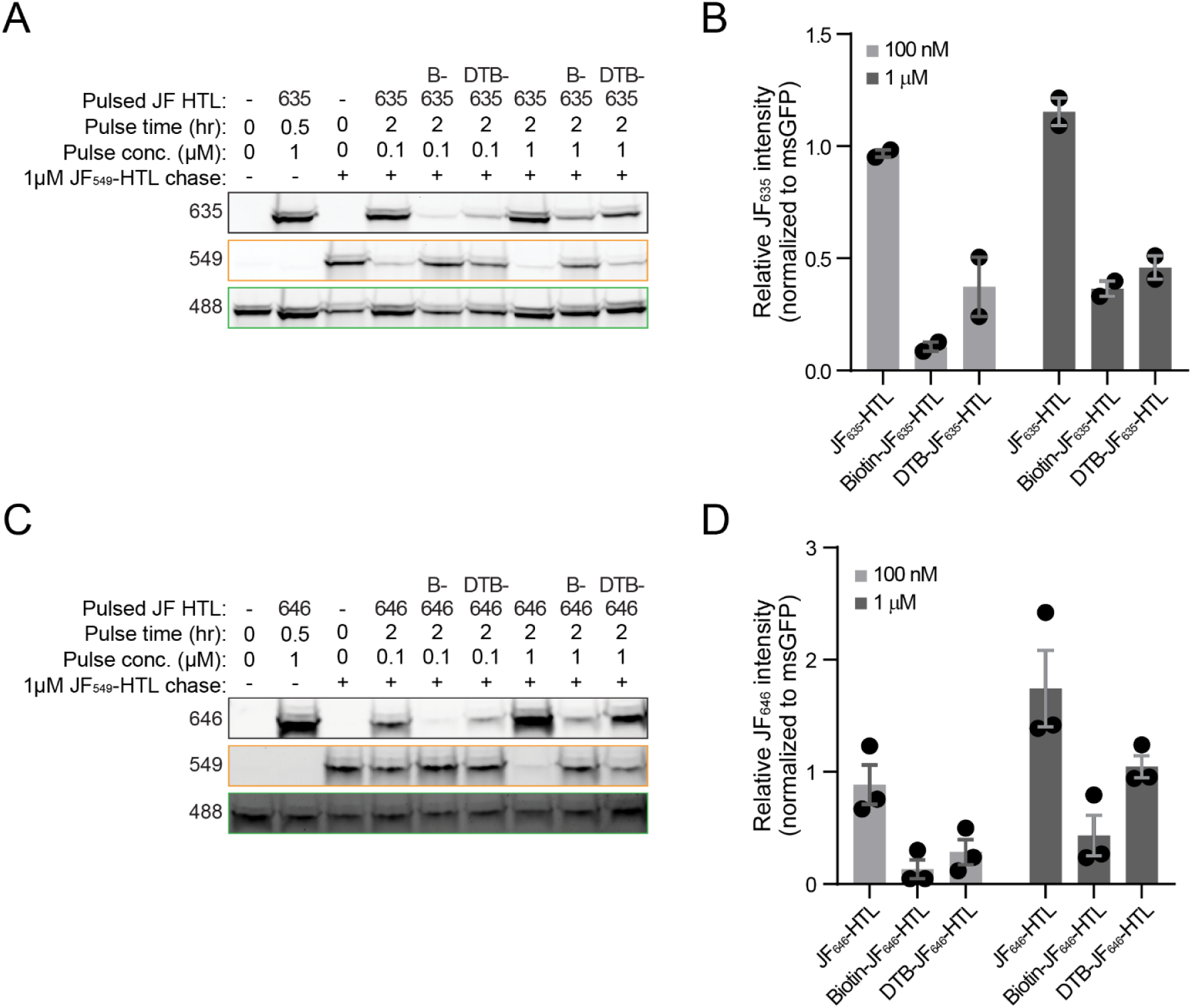
Permeability assay to assess intracellular availability of desthiobiotin-Janelia Fluor-HaloTag ligands (DTB-JF-HTLs). (**A**) In-gel fluorescence of SDS-PAGE gels for lysates collected after pulsing HEK293 cells expressing msGFP-HaloTag with a JF-HTL (JF_635-_HTL, biotin-JF_635-_HTL, or DTB-JF_635-_HTL) for 2 h at a concentration of 100 nM or 1 µM, and chasing with 1 µM JF549-HTL for 30 min, with the msGFP 488-nm signal shown as a loading control. (**B**) Quantification of gels shown in (A): 635-nm fluorescence corresponding to the JF-HTL pulse were normalized to the msGFP 488-nm signal. The graph depicts mean ± S.E.M. for two independent experiments. (**C**) In-gel fluorescence of SDS-PAGE gels for lysates collected after pulsing HEK293 cells expressing msGFP-HaloTag with a JF-HTL (JF_646-_HTL, biotin-JF_646-_HTL, or DTB-JF_646-_HTL) for 2 h at a concentration of 100 nM or 1 µM, and chasing with 1 µM JF_549-_HTL for 30 min, with the msGFP 488-nm signal shown as a loading control. (**D**) Quantification of gels shown in (C): 646-nm fluorescence corresponding to the JF-HTL pulse were normalized to the msGFP 488-nm signal. The graph depicts mean ± S.E.M. for three independent experiments.

**Extended Data 2.**
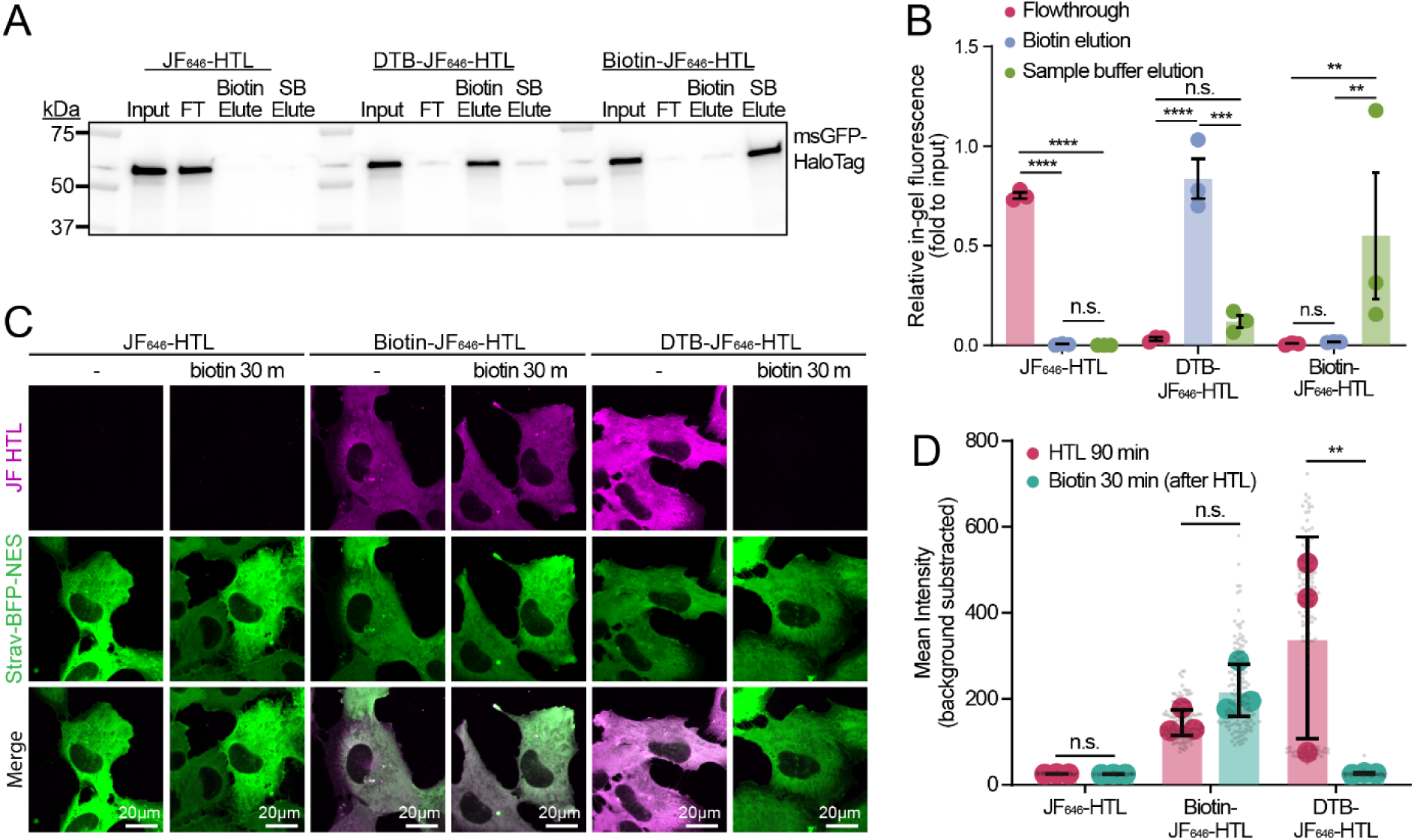
Assessing reversible binding of desthiobiotin-Janelia Fluor_646-_HaloTag ligand (DTB-JF_646-_HTL) and biotin-JF_646-_HTL to streptavidin. (**A**) In-gel fluorescence SDS-PAGE gel showing the results of a streptavidin pulldown assay comparing input, flowthrough (FT), the first biotin elution, and the second elution using sample buffer (SB) with heat for JF_646-_HTL, DTB-JF_646-_HTL, and biotin-JF_646-_HTL. (**B**) Quantification of in-gel fluorescence SDS-PAGE gels shown in (A) normalized to the input (n=3 independent experiments). The graph depicts mean ± S.E.M. Significance was tested using a two-way ANOVA with pairwise multiple comparisons; n.s., not significant; \*\**P* < 0.01; \*\*\**P* < 0.001; \*\*\*\**P* < 0.0001. (**C**) Confocal maximum intensity projection images of RPE1 cells expressing streptavidin-mtagBFP2-NES (green) which were incubated with JF_646-_HTL, biotin-JF_646-_HTL, or DTB-JF_646-_HTL to test their capacity to bind streptavidin. After a 2-h pulse, 80 µM biotin was added for 30 min to test if each HTL could be displaced. (**D**) Mean fluorescence intensity per cell (background subtracted) ± S.D. for HTL binding experiment (depicted in C) for three independent experiments (depicted by large, colored points) with n=100–149 cells/treatment (depicted by small, grey points). Significance was tested using a two-way ANOVA with pairwise multiple comparisons; n.s., not significant; \*\**P* < 0.01.

**Extended Data 3.**
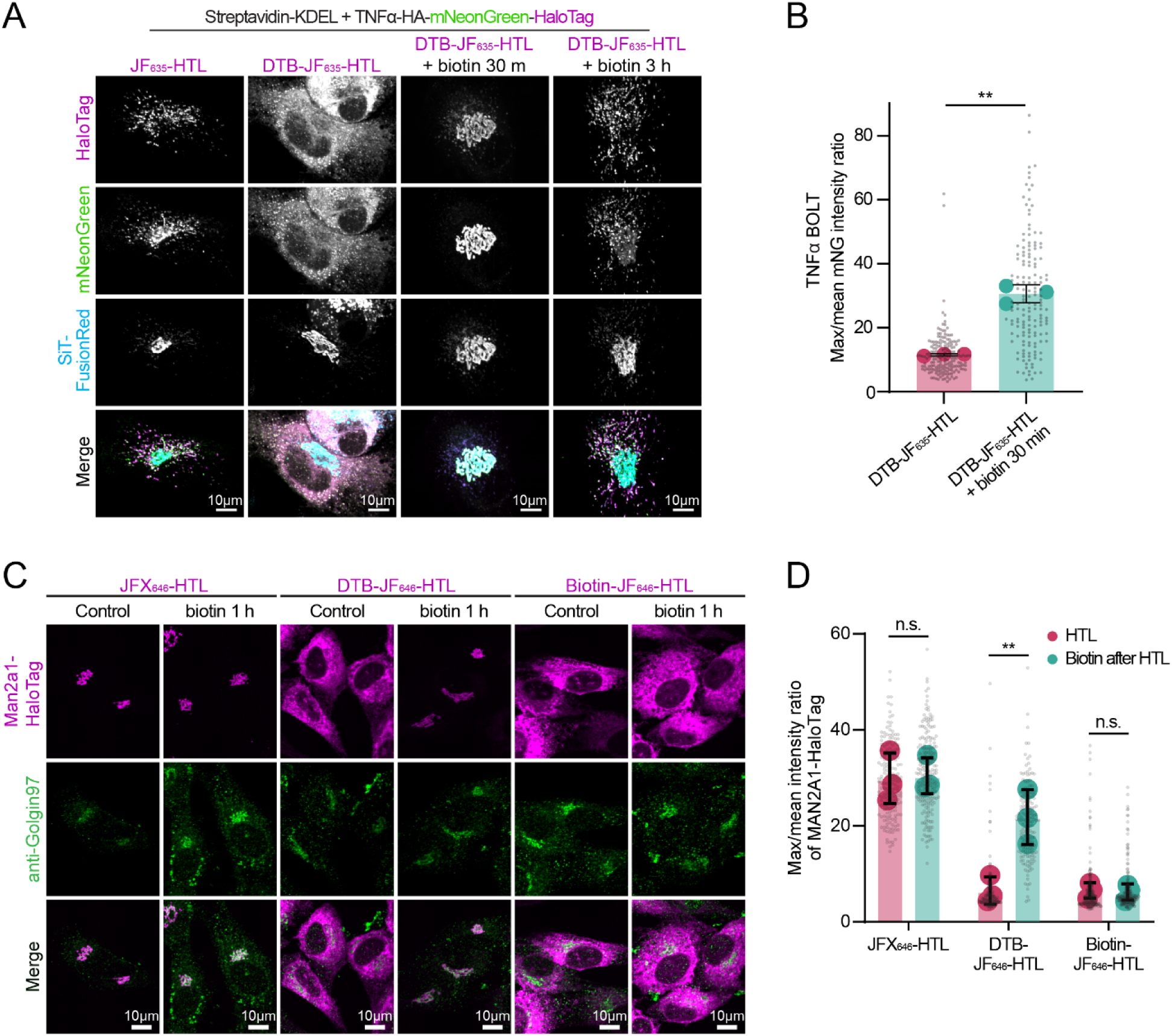
Applying BOLT to monitor endoplasmic reticulum (ER)-to-Golgi protein trafficking for TNFα-HA-mNeonGreen (mNG)-HaloTag synthetic cargo or endogenous MAN2A1-HaloTag knock-in RPE1 cells. (**A**) Confocal maximum intensity projections (MIPs) of RPE1 cells transduced with lentivirus encoding streptavidin-KDEL IRES TNFα-HA-mNG-HaloTag and SiT-FusionRed and incubated with 100 nM desthiobiotin-Janelia Fluor_635-_HaloTag ligand (DTB-JF_635-_HTL) or JF_635-_HTL overnight, then released via addition of biotin (80 µM final concentration) for 30 or 180 min. (**B**) The graph shows max-to-mean intensity ratio per cell for mNG fluorescence in images shown in (B), with the bars depicting the mean ± S.D. for three independent experiments (shown as large, colored points) n=158–179 total cells per treatment (shown as small, grey points). Statistics were performed using a paired *t-*test for three experimental means; ** *P* < 0.01. (**C**) Confocal MIPs of MAN2A1-HaloTag knock-in RPE1 cells transduced with lentivirus encoding streptavidin-KDEL and incubated with 100 nM JFX_646-_HTL, DTB-JF_646-_HTL, or biotin-JF_646-_HTL overnight to assess trapping in the ER and released with or without 80 µM biotin for 1 h to assess trafficking from ER to Golgi. Cells were fixed and immunostained for Golgin97 to label the Golgi. (D) Max-to-mean intensity ratio per cell for HTL fluorescence shown in (C), with the bars depicting the mean ± S.D. for three independent experiments (shown as large, colored points) with n=163–229 cells/condition (indicated as small, grey points). Statistics were analyzed using a two-way ANOVA with *post hoc* Sidak’s multiple comparisons testing. n.s., not significant; \*\**P* < 0.01.

**Extended Data 4.**
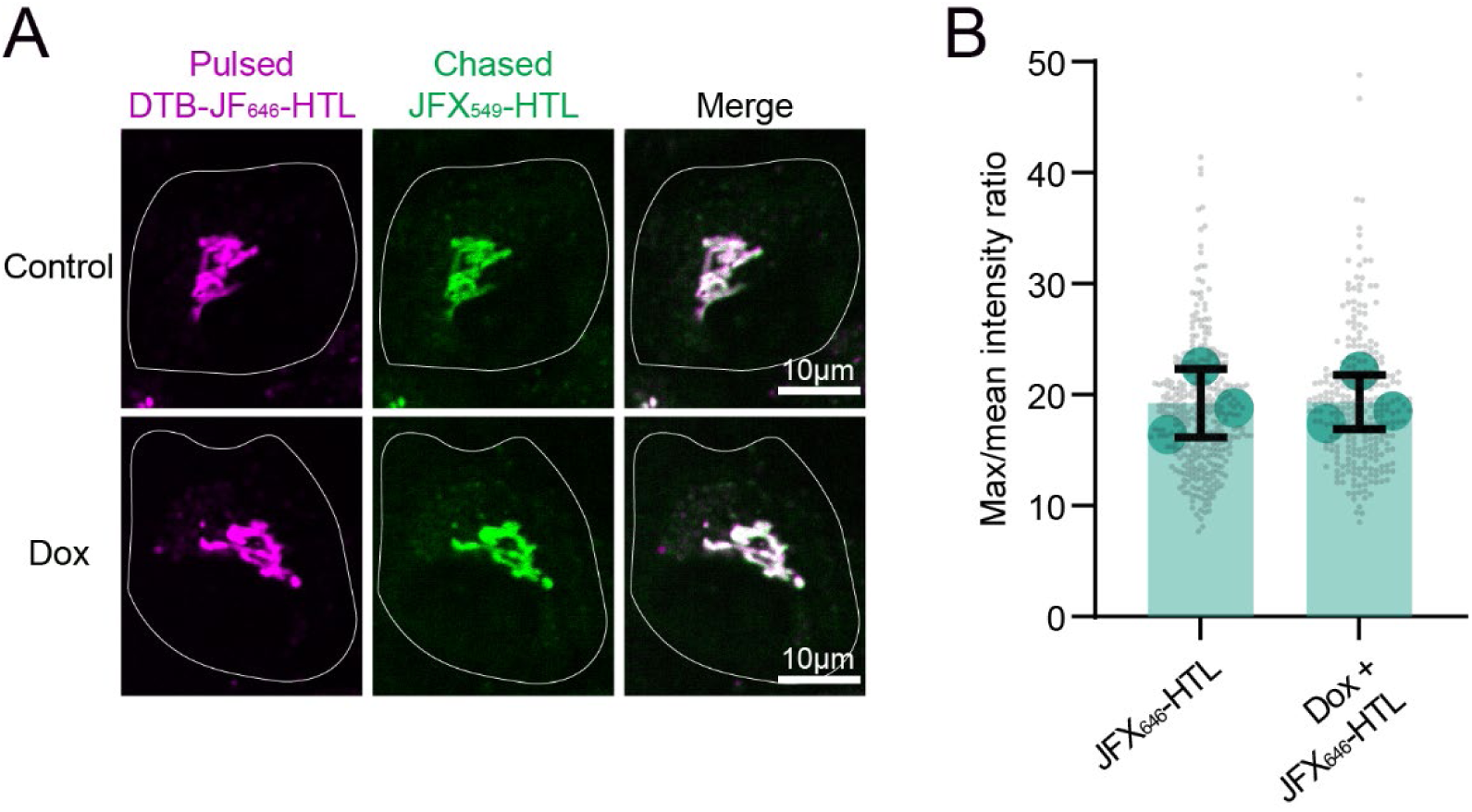
Negative control experiment showing JFX_646-_HTL lacking a desthiobiotin moiety cannot induce endogenous MAN2A1-HaloTag to recycle from the Golgi to the endoplasmic reticulum. (**A**) Confocal maximum intensity projections of MAN2A1-HaloTag knockin RPE1 cells that were pulsed overnight with 100 nM JFX646-HTL and chased with 100 nM JFX549-HTL for 6 h ± 1 µg/mL doxycycline (dox) to induce streptavidin-KDEL expression. (**B**) Max-to-mean intensity rations of images from (A). The graph depicts mean ± S.D. for three independent experiments (shown as large, colored points) with n=268–336 total cells per condition (indicated by small, grey points). A Welch’s *t*-test was performed to test statistical significance which was found to be not significant.

**Extended Data 5.**
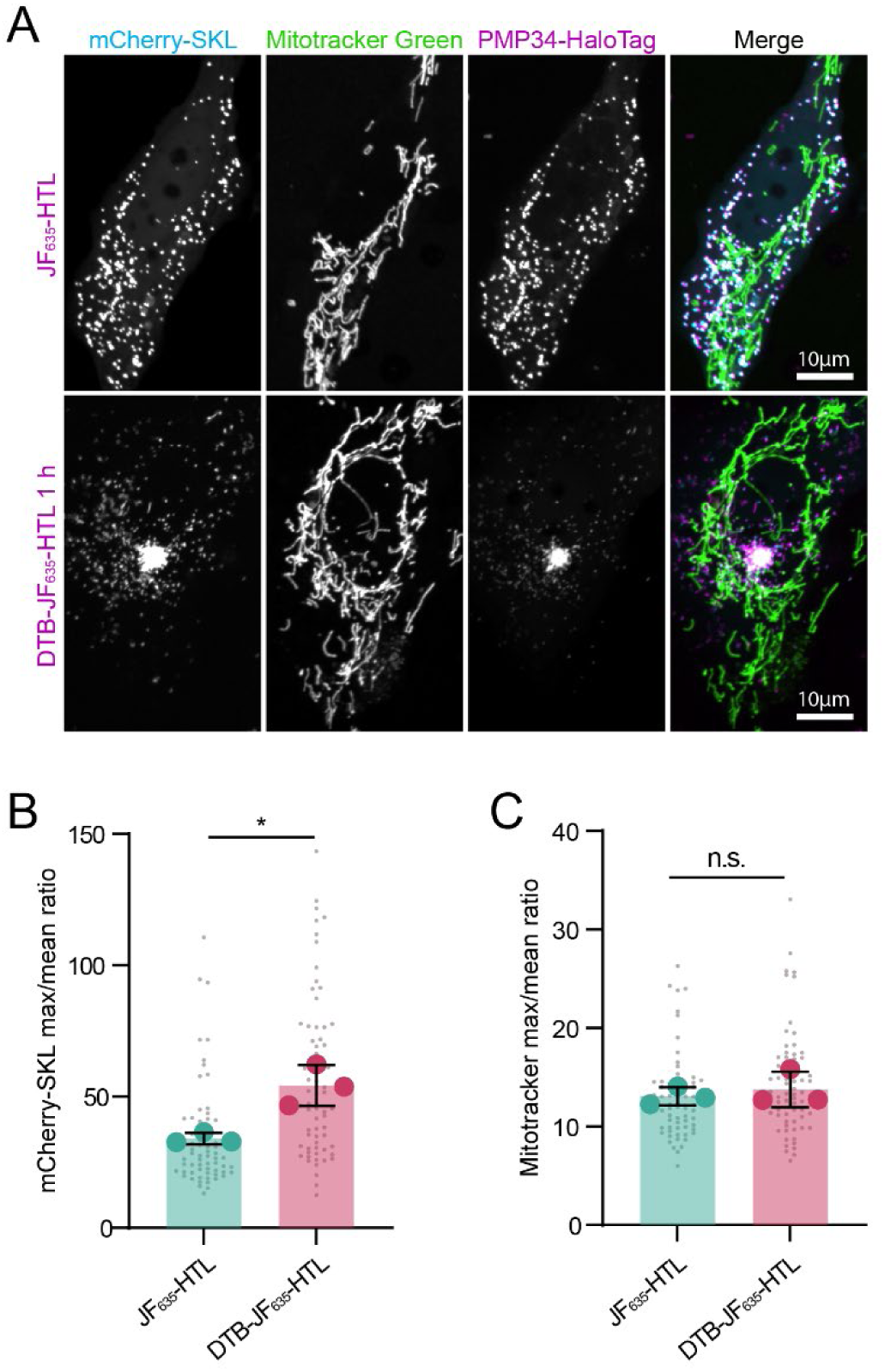
Trapping peroxisomes with PMP34-HaloTag BOLT does not trap mitochondria. (**A**) Confocal 2D images from timelapse imaging of RPE1 cells that were transduced with lentiviruses encoding flag-BicD2-streptavidin and mCherry-SKL, transfected with PMP34-HaloTag, and stained with MitoTracker Green FM. Cells were incubated with 100 nM JF_635-_HTL (negative control) or 100 nM DTB-JF_635-_HTL for 1 h to assess if mitochondria could co-traffic with peroxisomes when PMP34-HaloTag was induced to traffic peroxisomes to the cell center via BICD2-streptavidin recruitment. (**B**) Max-to-mean intensity ratios of mCherry-SKL and MitoTracker Green in images from (A). The graph depicts mean ± S.D for three independent experiments (shown as large, colored points) with n=64–68 cells/condition (shown as small, grey points). A Welch’s *t*-test was performed to test statistical significance. n.s., not significant; \**P* < 0.05.

