## Supplementary material for "Biotin-operated ligand trapping enables dynamic spatial redistribution of protein and organelle in living cells": Table S2

| Name | Sequence (5’ to 3’) | Description |
| --- | --- | --- |
| CAGE1043.MAN2A1.g6 | UCAAUCCAAAUGUGAAAGUC | gRNA spacer |
| CAGE1043.MAN2A1.DS.F | CTACACGACGCTCTTCCGATCTtcgaaagtctcacaccttcatc | Primer flanking cut site |
| CAGE1043.MAN2A1.DS.R | CAGACGTGTGCTCTTCCGATCTttctcacagtagccagaagctaa | Primer flanking cut site |
| CAGE1043.halo.5gen.DS.F | CTACACGACGCTCTTCCGATCTactgtatttgcactctattgtattgttga | 5’ junction primer |
| CAGE1043.halo.5junc.DS.R | CAGACGTGTGCTCTTCCGATCTaaagccagtaccgatttctgccatgcc | 5’ junction primer |
| CAGE1043.halo.3junc.DS.F | CTACACGACGCTCTTCCGATCTtcgacgctcgagatttccggcatgtac | 3’ junction primer |
| CAGE1043.halo.3gen.DS.R | CAGACGTGTGCTCTTCCGATCTaggcacatttattttgatatggt | 3’ junction primer |
| CAGE1043.g6.Halo.dsDNA | gatctaagtttacttggttctgagcattctgacattaaggccttgagtcataagggggttatacgatccagagagacagataacatggcagcggtgagggaaatactttccaaatgacctgtccacctctatagactgggactcaatgacataatgatccctccttgtagtgagcctgtgtgctctctcttaactctgttggaacaggaaaaagtggttgaggactaccattgttaggggtaagtattgactaaaagagaatgaggatgtttttaaagcttcttggccattggaggtcctttcttttattgaaaaagcccctgtgcatgataaaattctcaagcaatacaaaaaggaatgcagaatatgttgctcatgaccattactggtttcctgttcccttccaagcgtagtctgtatgttcaagcatgtatatatgtatatttaattcacccttttttctgtaaaagagaacttatatatgtatatttaataacatatatgtatatttaatccacccttttttcctcaaaagagaacttcaacatactgttttattccttctcatactaattttcacagaatatgtagttgtgctgctaatcttcaaagttgatctaaatatgtaaattttatcatttacttttcagatattggtacagaaacttttaaacaagtttattgtcgaaagtctcacaccttcatcactatccttgatgcattcacctcccggcactcagaatataagtgagatcaacttgagtccaatggaaatcagcacattccgaatccagttgaggtcagctggaggcagtgcaggtggcatggcagaaatcggtactggctttccattcgacccccattatgtggaagtcctgggcgagcgcatgcactacgtcgatgttggtccgcgcgatggcacccctgtgctgttcctgcacggtaacccgacctcctcctacgtgtggcgcaacatcatcccgcatgttgcaccgacccatcgctgcattgctccagacctgatcggtatgggcaaatccgacaaaccagacctgggttatttcttcgacgaccacgtccgcttcatggatgccttcatcgaagccctgggtctggaagaggtcgtcctggtcattcacgactggggctccgctctgggtttccactgggccaagcgcaatccagagcgcgtcaaaggtattgcatttatggagttcatccgccctatcccgacctgggacgaatggccagaatttgcccgcgagaccttccaggccttccgcaccaccgacgtcggccgcaagctgatcatcgatcagaacgtttttatcgagggtacgctgccgatgggtgtcgtccgcccgctgactgaagtcgagatggaccattaccgcgagccgttcctgaatcctgttgaccgcgagccactgtggcgcttcccaaacgagctgccaatcgccggtgagccagcgaacatcgtcgcgctggtcgaagaatacatggactggctgcaccagtcccctgtcccgaagctgctgttctggggcaccccaggcgttctgatcccaccggccgaagccgctcgcctggccaaaagcctgcctaactgcaaggctgtggacatcggcccgggtctgaatctgctgcaagaagacaacccggacctgatcggcagcgagatcgcgcgctggctgtcgacgctcgagatttccggcatgtacaagtaaaactgactttcacatttggattgagaatcattggcttttatacctttcttggtttgacgtgcaataaagaagcacattattttagcttctggctactgtgagaacatgaattctgtgattctgtgggttttttcttttttcttttaccagtacagtaagaaaaaaaaaaaaaaaaaaaaagccatgctatcaatcaagattctttttttttaaactttctcccatgaactaccaccatcagtatgaattgatgcaacaaatgaagaaatatttaaagacagcctctcaacagattgtatctcaggttaaatgctaactaattatgtctgtgttgggggttgcgaagagattcttaaaagtatctgtgtgttgatcatcagttttacaaaaacacctatttggctgaaatggaataaaatgtttgtgggtaaaagctaatggccaaaatggttgcaatcattcatactagttagaaaaattatgtgttgaaataagtggaaaagtgcaatccatccacccttatgattaacgtagatgatttttatacctttttctgatgtacctcttgaccttctccttcccttcctaccctttctaagtatttccagaaatacctgattttgaatcattcaacagtagaaaaagaggcatattttcattacttgacaatgtgggatgggtgcaatttattccatcttcactaaaatagaagcaattccataggtaccataaacctattttaggtaccacaaggtgtctttttacacagctcatttgaatacaggtgttctgagaaggggtttctat | dsDNA donor  pUC57 backbone not shown |
| CAGE1046.LMAN1.g30 | CUCCUCCCACGCCGUCGCCC | gRNA spacer |
| CAGE1046.LMAN1.DS.F | CTACACGACGCTCTTCCGATCTctgccaatcagcgagcgccctgttg | Primer flanking cut site |
| CAGE1046.LMAN1.DS.R | CAGACGTGTGCTCTTCCGATCTgcgaagaggcagcagaagggggaga | Primer flanking cut site |
| CAGE1046.LMAN1.5gen.DS.F | CTACACGACGCTCTTCCGATCTtgtagcccacacgaacagag | 5’ junction primer |
| CAGE1046.LMAN1.5junc.DS.R | CAGACGTGTGCTCTTCCGATCTacttgtggccgtttacgtcg | 5’ junction primer |
| CAGE1046.LMAN1.3Junc.DS.F2 | CTACACGACGCTCTTCCGATCTtcgccgaccactaccagcag | 3’ junction primer |
| CAGE1046.LMAN1.3gen.DS.R | CAGACGTGTGCTCTTCCGATCTtgccaagtgctacgattgact | 3’ junction primer |
| CAGE1046.LMAN1.dsDNA | ctctcaccaaaaccaaaccaaaccaaagtcagtttcatttgaatgcactcttgccatggtttcaaaatttggaaagctcttggaattaaattttgagatgtaaaggctggctgacaaattcacagaaaccctctttccgacaccaacttataaatcctaaggcatcctacaaccccgtgggtgctggtttctagtttgcaacgctctaacatagcttggtagagctcagtgggctgggaccataagcattcgggattgtgccactgaaaaactgatgtcacatcaaaaggaacacctttcttctaatatactaaaaaggttttgtacaggctgggtccagaggtcttagggctagccaattcctgctcccaggtattctaagaagccggagaaagctccctgcaggtaaggacccatcggcgcttcccgcactctgccgcgcgggaagcgaaggttggagtcctttccccacccccggagggtgggcggggccgaaggaccgcaccgccccagacccgccccctgcgcgtggccccgcccacagctgctcctggccgtgcggccgcgcggaaagcggaggccgggggcggggctcaccccttgcccttgggggtcccccgggctctgccaatcagcgagcgccctgttggccatccgcagccatccgcgcccctccctctcccgccctcctccgcgttccagaatccaagatggcgggatccaggcaaaggggtctccgggccagagttcggccgctgttctgcgccttgctgctgtcactcggtcgcttcgtccggggcATGGTGAGCAAGGGCGAGGAGCTGTTCACCGGGGTGGTGCCCATCCTGGTCGAGCTGGACGGCGACGTAAACGGCCACAAGTTCAGCGTGTCCGGCGAGGGCGAGGGCGATGCCACCTACGGCAAGCTGACCCTGAAGTTCATCTGCACCACCGGCAAGCTGCCCGTGCCCTGGCCCACCCTCGTGACCACCTTGACCTACGGCGTGCAGTGCTTCGCCCGCTACCCCGACCACATGAAGCAGCACGACTTCTTCAAGTCCGCCATGCCCGAAGGCTACGTCCAGGAGCGCACCATCTTCTTCAAGGACGACGGCAACTACAAGACCCGCGCCGAGGTGAAGTTCGAGGGCGACACCCTGGTGAACCGCATCGAGCTGAAGGGCATCGACTTCAAGGAGGACGGCAACATCCTGGGGCACAAGCTGGAGTACAACTACAACAGCCACAAGGTCTATATCACCGCCGACAAGCAGAAGAACGGCATCAAGGTGAACTTCAAGACCCGCCACAACATCGAGGACGGCAGCGTGCAGCTCGCCGACCACTACCAGCAGAACACCCCCATCGGCGACGGCCCCGTGCTGCTGCCCGACAACCACTACCTGAGCACCCAGTCCAAGCTGAGCAAAGACCCCAACGAGAAGCGCGATCACATGGTCCTGCTGGAGTTCGTGACCGCCGCCGGGATCACTCTCGGCATGGACGAGCTGTACAAGTCAGCTGGAGGCAGTGCAGGTGGCgacggcgtgggaggagaccccgcggtcgcgttgccacatcgccgtttcgagtacaaatacagcttcaaggggccgcacctggtgcagagcgacgggaccgtgcccttctgggcccacgcggggagtaagccggagcagagggctgggggccgggttcctctcccccttctgctgcctcttcgcctttcatccgcggctaccctggtgtgctgaggttcccgttccagcagctgcacaccatatgcgctgtggaggggaccctcctggcagcccctccgccgggtccctcctggttcgcacggttccctccgcttgtccccgagtctttccctcacccattctgttctctctcacccgcctgctgtcttcccttctttgatctcgtcggtactttccgttggatctagaggtgtttggttacctcctacacttcctagcactcaaacttgcttctaggtatcatgtttcacaacgccacgtctctaggagaaatgtgccctacccccagccatcccaacctgagacatgttctagaagcccccttacttcctttcccgcttccaccccccacccccgatccccgatttctttgctccggtgttagcaatggttaatgcttcgcttgaaccccatgcccactgacggctcctgagatggcatatagtagtattagagtaattttttaagtctttgtatgtcagaagacattcttatactgttggaactacacattttcctctcagtgacttggtgtggtctttgatcttttctctcgtagtaacacattatgtaacttttta | dsDNA donor  pUC57 backbone not shown |
