## Supplementary material for "Biotin-operated ligand trapping enables dynamic spatial redistribution of protein and organelle in living cells": Table S3

**Table S3.** Sequences of gRNA and primers used for knockin and validation of TP53-HaloTag-V5 cell line.

| Oligos | Sequence (5’ to 3’) |
| --- | --- |
| TP53 gRNA sequence | GGAGAATGTCAGTCTGAGTC |
| TP53 Junction primer | TATTTGCATGGGGTGTGGCC |
| HaloTag Junction primer | ATGATGTTGCGCCACACGTAGG |
| HaloTag Sanger primer | CTCGCCCAGGACTTCCACAT |
